# The Norwegian Mother, Father, and Child cohort study (MoBa) genotyping data resource: MoBaPsychGen pipeline v.1

**DOI:** 10.1101/2022.06.23.496289

**Authors:** Elizabeth C. Corfield, Alexey A. Shadrin, Oleksandr Frei, Zillur Rahman, Aihua Lin, Lavinia Athanasiu, Bayram Cevdet Akdeniz, Tahir Tekin Filiz, Laurie Hannigan, Robyn E. Wootton, Chloe Austerberry, Amanda Hughes, Martin Tesli, Lars T. Westlye, Hreinn Stefánsson, Kári Stefánsson, Pål R. Njølstad, Per Magnus, Neil M. Davies, Vivek Appadurai, Gibran Hemani, Eivind Hovig, Tetyana Zayats, Helga Ask, Ted Reichborn-Kjennerud, Ole A. Andreassen, Alexandra Havdahl

## Abstract

**Background:** The Norwegian Mother, Father, and Child Cohort Study (MoBa) is a population-based pregnancy cohort, which includes approximately 114,500 children, 95,200 mothers, and 75,200 fathers.

Genotyping of MoBa has been conducted through multiple research projects, spanning several years; using varying selection criteria, genotyping arrays, and genotyping centres. MoBa contains numerous interrelated families, which necessitated the implementation of a family-based quality control (QC) pipeline that verifies and accounts for diverse types of relatedness.

**Methods:** The MoBaPsychGen pipeline, comprising pre-imputation QC, phasing, imputation, and post-imputation QC, was developed based on current best-practice protocols and implemented to account for the complex structure of the MoBa genotype data. The pipeline includes QC on both single nucleotide polymorphism (SNP) and individual level. Phasing and imputation were performed using the publicly available Haplotype Reference Consortium release 1.1 panel as a reference. Information from the Medical Birth Registry of Norway and MoBa questionnaires were used to identify biological sex, year of birth, reported parent-offspring (PO) relationships, and multiple births (only available in the offspring generation).

**Results:** In total, 207,569 unique individuals (90% of the unique individuals included in the study) and 6,981,748 autosomal SNPs passed the MoBaPsychGen pipeline. A further 174,462 chromosome X and 3,200 PAR SNPs are available in a subset of these individuals (N = 204,913 and 135,593, respectively). The relatedness checks performed throughout the pipeline allowed identification of within-generation and across-generation first-degree, second-degree, and third-degree relatives. The individuals passing post-imputation QC comprised 64,471 families ranging in size from singletons to 84 unique individuals (singletons are included as families as other family members may not have been genotyped, imputed, or passed post-imputation QC). The relationships identified include 287 monozygotic twin pairs, 22,884 full siblings, 117,004 PO pairs, 23,299 second-degree relative pairs, and 10,828 third-degree relative pairs.

**Discussion:** MoBa contains a highly complex relatedness structure, with a variety of family structures including singletons, PO duos, full (mother, father, child) PO trios, nuclear families, blended families, and extended families. The availability of robustly quality-controlled genetic data for such a large cohort with a unique extended family structure will allow many novel research questions to be addressed. Furthermore, the MoBaPsychGen pipeline has potential utility in similar cohorts.

## Introduction

In 2005, the first genome-wide association (GWA) study was published [1]. Since then, the field has continued to advance with GWA studies of ever-increasing sample sizes and imputation with improved reference panels. This has allowed the identification of thousands of single nucleotide polymorphisms (SNPs) associated with a multitude of human traits [2]. Advancements in statistical methodologies and modelling software have allowed increasingly complex research questions to be addressed, for example, using multivariate GWA analysis [3], polygenic score analysis [4], and Mendelian Randomization [5].

As the field has progressed, scripted open-source quality control (QC) and imputation pipelines for genotype data, such as the Rapid Imputation and COmputational PIpeLIne for Genome-Wide Association Studies (RICOPILI) [6], have been developed. These pipelines are designed to be arbitrarily used by cohorts or studies and are typically updated to include the most up-to-date best-practice procedures [7, 8]. Despite continued improvements in QC practices, handling complex relatedness in processing of genotyped samples remains a major challenge.

Many of the largest longitudinally studied pregnancy and birth cohorts available today consist of related individuals. A key benefit of cohorts with related individuals is the ability to perform complex genetic analyses, by explicitly modelling the relatedness. Cohorts with large numbers of related individuals can be used to address unique research questions, which would not be possible in cohorts comprising only unrelated individuals. These include investigations of genetic and environmental mechanisms of intergenerational transmission [9], and interrogations of the role of familial confounding in observational associations [10].

However, to exploit the novel analytical opportunities available using increasingly complex family cohorts, family-based QC protocols needs to be performed, otherwise the researcher risks inducing a bias in results. To our knowledge, the Pedigree Imputation Consortium Pipeline (PICOPILI) [11] is the most used scripted open-source family-based QC pipeline available. PICOPILI offers scripts for pre-imputation QC, phasing, imputation, post-imputation QC, and GWA of family data with arbitrary pedigrees, alongside tools for imputation and family-based analyses. However, some of the software used in PICOPILI is not designed to handle large sample sizes. Furthermore, generic open-source pipelines such as PICOPILI are – by design – not equipped to handle idiosyncrasies of specific cohorts or studies.

The Norwegian Mother, Father, and Child Cohort Study (MoBa) is a population-based pregnancy cohort with a complex relatedness structure and multiple ancestries [12]. The cohort contains several types of relationships, including monozygotic (MZ) and dizygotic (DZ) twin pairs, parent-offspring (PO) duos and trios, full-siblings (FS), half-siblings (HS), aunt/uncle-niece/nephew (AUNN) and grandparent-grandchild (GO) pairs, and first cousins. Additionally, genotyping in MoBa was not conducted as a single systematic project, which has resulted in genotyping batches with varying sample sizes, selection criteria, and genotyping arrays. The considerable number of variously interrelated families in MoBa combined with the genotyping complexities necessitates the implementation of a family-based QC pipeline that verifies and accounts for diverse types of relatedness, while appropriately handling the differences resulting from a varied genotyping strategy, such as assessment of genotyping plate and genotyping batch effects.

Here, we report in full the design and implementation of the MoBaPsychGen genotype QC pipeline: a novel family-based pipeline, which includes pre-imputation QC, phasing, imputation, and post-imputation QC. We provide details of data resulting from the implementation of this pipeline in the MoBa cohort, including the number and nature of familial relationships among individuals passing QC.

## Materials and Methods

### MoBa

MoBa is a population-based pregnancy cohort study conducted by the Norwegian Institute of Public Health. Participants were recruited from all over Norway from 1999 to 2008. The women consented to participation in 41% of the pregnancies (N = 112,908 recruited pregnancies) [13]. The cohort includes approximately 114,500 children, 95,200 mothers and 75,200 fathers. Blood samples were obtained from both parents during pregnancy and from mothers and children (umbilical cord) at birth [14]. The current study is based on version 12 of the quality-assured data files released for research in January 2019.

The establishment of MoBa and initial data collection was based on a license granted from the Norwegian Data Protection Agency and an approval from The Regional Committees for Medical and Health Research Ethics (REK). The MoBa cohort is currently regulated by the Norwegian Health Registry Act. The current study was approved by The Regional Committees for Medical and Health Research Ethics (14140 and 2016/1226). In accordance with REK regulations, individuals who withdraw consent are excluded.

### Reported pedigree and sex information

Information from the Medical Birth Registry of Norway (MBRN), a national health registry containing information about all births in Norway, and the MoBa questionnaire data were used to identify sex, year of birth, multiple births (in the offspring generation), and reported PO relationships. Prior to genetic analyses, all pedigrees were constructed based on the reported PO relationships for each pregnancy. Each family contained all reported relationships (these included: PO, FS from pregnancies with single and multiple births, and HS relationships). Wherever possible, sex was assigned using information from the MBRN. In instances where sex was not specified in MBRN, the reported sex from the MoBa questionnaires was used.

### Biological data

Blood samples were collected from participating mothers and fathers at approximately the 17^th^ week of pregnancy during the ultrasound examination. A second blood sample was taken from the mother soon after birth. The blood sample for the child was taken from the umbilical cord after birth. Biological samples were sent to the Norwegian Institute of Public Health where DNA was extracted by standard methods and stored [15].

### Genotyping

Genotyping of MoBa has been conducted through multiple research projects, spanning several years. The research projects (HARVEST, SELECTIONpreDISPOSED, and NORMENT) provided genotype data to MoBa Genetics (https://github.com/folkehelseinstituttet/mobagen). In total, 238,001 MoBa samples were sent to be genotyped in 24 genotyping batches with varying selection criteria, genotyping arrays, and genotyping centres (Supplementary Table 1). Two genotyping batches were split into sub-batches for technical reasons; the 26 batches and sub-batches are henceforth referred to as batches. Full details about genotyping of MoBa (including details about the sub-batches) are described in the supplementary text (eMethods1).

As a quality control measure, 397 individuals were deliberately genotyped more than once. Furthermore, genotyping efforts were conducted independently, which resulted in numerous individuals being genotyped multiple times. In total, there were 4,035 individuals genotyped multiple times, of which 3,958 were genotyped twice and 77 genotyped three times.

### Implementation of the family based MoBaPsychGen genotype QC pipeline

The MoBaPsychGen pipeline for QC and imputation was developed to ensure the complex relationship structure and varying selection criteria, genotyping batches, and genotyping arrays were handled appropriately. QC was performed based on current best-practice protocols [6-8, 11, 16]. Throughout the pipeline, both SNP and individual level QC were performed; with priority given to retaining individuals over SNPs, as SNPs can be imputed. The complete pipeline is described in text module-by-module below and a simplified overview of the pipeline is shown in Figure 1. The primary software used throughout the pipeline for QC was PLINK1.9 [17] and The R Project for Statistical Computing [18] was used to produce figures. Further explanation of specific decisions relating to thresholds or software utilized is included in the supplementary text. Scripts used throughout the MoBaPsychGen pipeline are available on GitHub: https://github.com/psychgen/MoBaPsychGen-QC-pipeline.

**Figure 1.**
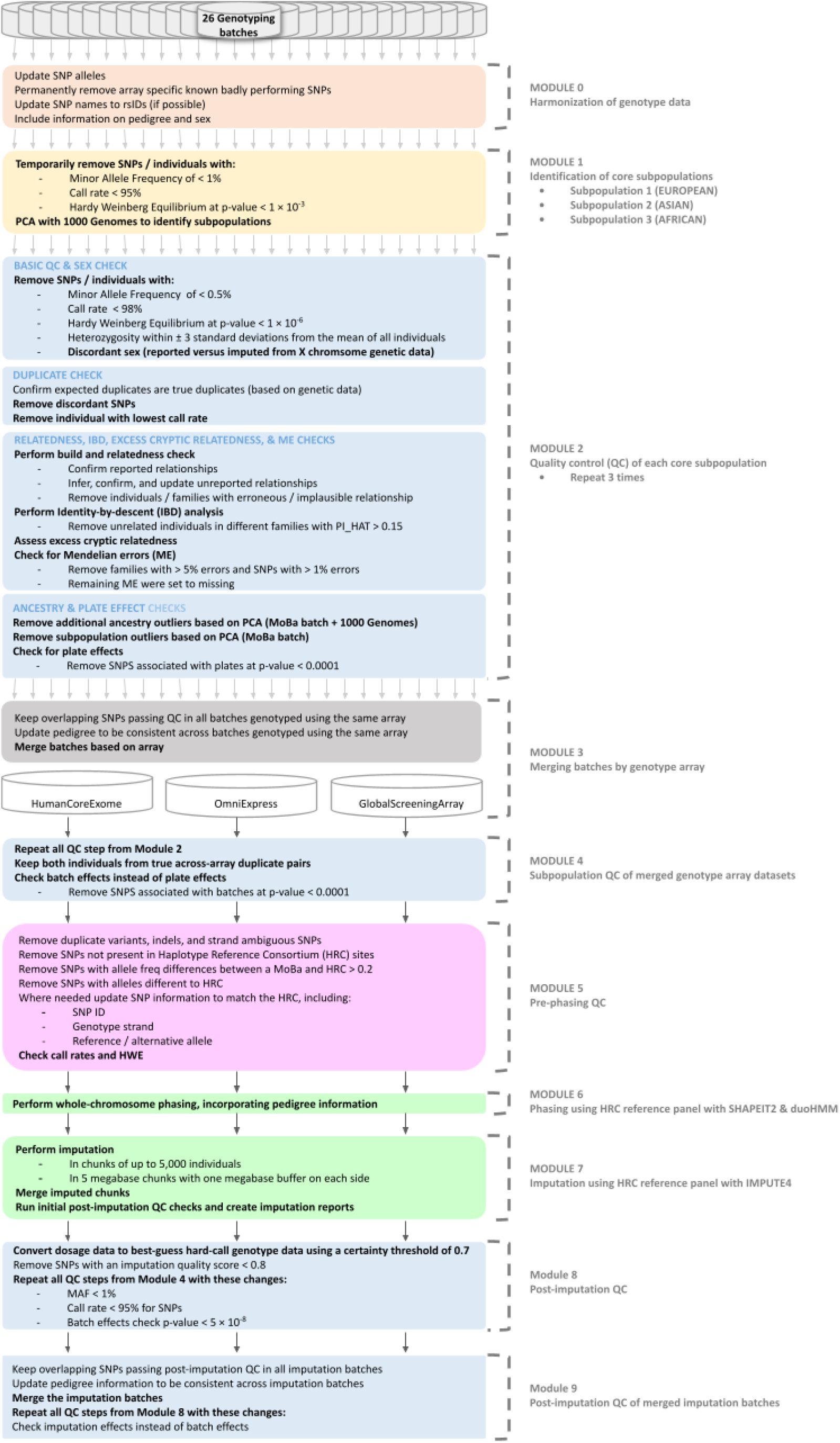
Simplified flowchart of the pipeline. Figure abbreviations are SNP = single-nucleotide polymorphism, PCA = principal component analysis, and PI_HAT = proportion of the genome shared identity-by-descent.

### Module 0: Harmonize genotype data

Initially, harmonization of the data was conducted (as needed for each batch), including (1) updating SNP alleles (convert from Illumina A/B alleles to the genetic A, C, T, and G alleles); (2) excluding known problematic SNPs previously reported by the Psychiatric Genomics Consortium (PGC) [19]; (3) updating SNP names to rsIDs; and (4) converting the chromosomal positions to match the genome build of the phasing and imputation reference panel (GRCh37/hg19), using the LiftOver tool [20]. The genotype files (PLINK bfile format [21]) were then updated to include the reported pedigree and sex information.

### Module 1: Identify subpopulations

Initial pre-imputation QC was performed (using PLINK1.9 [17]) to identify subpopulations, as population stratification (systematic differences in minor allele frequency (MAF) between populations) can lead to spurious associations and needs to be accounted for during QC [22], however, the software currently available lack the methodologies to perform QC in admixed populations. SNPs were temporarily removed based on the following criteria: (1) call rate < 95%; (2) out of Hardy-Weinberg equilibrium (HWE) at p < 1.00 × 10^−3^; and (3) MAF < 1% (permanent SNP removal for call rate, HWE and MAF are performed in module 2). Individuals were temporarily removed with call rate ≤ 95% (permanent individual removal for call rate is performed in module 2). Principal component analysis (PCA) was then performed with 1000 Genomes phase 1 [23] (N = 1,083 unrelated individuals) to identify the subpopulations. To perform the PCA, linkage disequilibrium (LD) pruning (using the PLINK indep-pairwise parameter with a window size of 3000, step size of 1500, and r^2^ threshold of 0.1) was performed and SNPs in long-range high LD [24] regions were removed before merging the batch genotypes with those of the 1000 Genomes data [23]. Principal components (PC) were first estimated within founders (based on reported information). The non-founders, all individuals with at least one reported parent, were then projected into the PC space of founders. Individuals were then assigned to European, Asian, and African core subpopulations based on visual inspection using the first seven PCs.

### Module 2: QC of subpopulations

Pre-imputation QC of the subpopulations was performed (using PLINK1.9 [17]) in multiple rounds to ensure the validity of checks performed as individuals and SNPs were removed throughout the QC. For MoBa genotyping data, three rounds were enough to ensure a robust QC was performed with only SNPs and individuals of high quality passing through the pipeline. Module 2 was performed separately for each subpopulation as currently there is a lack of methodology to perform QC in admixed populations (for example heterozygosity cannot be estimated properly in admixed populations).

#### Basic QC

Initially a basic QC was performed where SNPs and individuals were removed based on the following filters: (1) SNPs with MAF < 0.5%; (2) SNPs with call rate < 95%; (3) SNPs with call rate < 98% (eMethods2); (4) SNPs and individuals with a call rate < 98%; and (5) SNPs out of HWE at p < 1.00 × 10^−6^. Heterozygosity filtering was then performed with individual outliers removed if they were ± 3 standard deviations from the mean heterozygosity across all individuals using only autosomes (eMethods3).

#### Sex check

A sex check was performed, including the following steps: (1) ensuring the pseudo-autosomal region is coded as a separate XY chromosome; (2) check sex assignments reported at birth against those imputed from X chromosome genotype calls; (3) identify individuals with problematic sex assignment—females with inbreeding coefficient (F) > 0.2 and males with F < 0.8; and (4) remove individuals with discordant sex—females with F > 0.5 and males with F < 0.5.

#### Duplicate QC

Duplicate QC was then performed, including the following steps: (1) confirming expected duplicates (based on phenotype information) are true duplicates, based on genetic data using PLINK [17] identity-by-descent (IBD) analysis with an estimated proportion of the genome shared IBD (PI_HAT) > 0.98; (2) removing all discordant non-missing SNPs, based on PLINK SNP concordance analysis in true duplicates; and (3) removing the individual with the lowest call rate from each true duplicate pair. Expected duplicates that were not confirmed in the IBD analysis were resolved in the relatedness check.

#### Relatedness checks

A pedigree build and relatedness check was performed using KING version 2.2.5 [25] (eMethods4), whereby the genetic data was used to confirm reported relationships and infer unreported relationships. In the presence of admixture, KING [25] accurately infers MZ twin or duplicate pairs (kinship coefficient > 0.3540), first-degree (PO, FS, DZ twin pairs; kinship coefficient range 0.1770 - 0.3540), second-degree (HS, GO, AUNN; kinship coefficient range 0.0884 – 0.1770), and third-degree (first cousins; kinship coefficient range 0.0442 – 0.0884) relationships. Any within-family errors or between-family issues that were identified in the relatedness check were investigated. All plausible identified relationships inferred from the genetic data were updated in the pedigree. Meanwhile, individuals or families were removed where needed, if the reported or inferred relationship was erroneous or implausible (for example being an unexpected duplicate). An in-depth description of how each relationship type was confirmed is described below.

Inferred MZ twin pairs were confirmed based on whether they had the same sex and year of birth (YOB). Furthermore, the relationships of identified MZ twin pairs with other family members were also examined to ensure they were as expected. When a pair of individuals were inferred as a duplicate/MZ twin pair, and reported information indicated the pair were unexpected duplicates (based on reported information it was impossible for the pair to be an MZ twin pairs, i.e., different sex, YOB, parents), the inferred family relationships were used to identify the individual to exclude from the study. Within-generation PO relationships and across-generation FS or HS relationships were confirmed together. The within-generation PO relationship was initially confirmed based on an age difference of at least 15 years, followed by confirmation that all other family relationships were as expected, including the across-generation FS or HS relationship. If the relationships could not be confirmed, unexpected family relationships were used to identify the individual(s) to remove. FS in parent generation were incorporated if all relationships between the families were as expected. FS and HS in the child generation were incorporated if the reported PO relationships were as expected and one or both parents were unreported. Inferred PO relationships where the parent was not reported were confirmed if all other expected relationships were as expected. When PO errors occurred where the expected mother was unrelated to the child, the family was removed as blood was taken from the mother and child at the same time. Meanwhile, for PO errors where an expected father was unrelated to the child, the PO relationship was broken and both individuals were kept in the family. The individuals were not removed or placed into separate families, because it is plausible (in contrast to unrelated mothers, described above) that the individual from whom the blood sample was collected as the father was not the biological father of the child but is acting as parental figure for the child, therefore, it is important to keep the individual in the dataset for gene-environment studies.

PLINK [17] IBD analysis was used to confirm the relationships inferred by KING [25]. Unrelated individuals who were not members of a family and had an estimated PI_HAT value greater than 0.15 were removed (approximately corresponding to the lower threshold for coding second-degree relatives or the upper threshold for coding third-degree relatives in KING).

An excess cryptic relatedness check was performed by, (1) computing a sum of kinship coefficients > 2.5%; (2) plotting the summed kinship for each individual; and (3) removing outliers based on visual inspection.

After within-family and between-family relationships were confirmed by genetic data, a check for Mendelian errors (ME) was performed in PLINK. The ME check included families with one or two parents present in the data. Families with > 5% errors and SNPs with > 1% errors were removed. The remaining ME were set to missing.

#### Ancestry and plate effect checks

Within the core subpopulations identified in module 1, additional ancestry outliers were removed based on PCA with the 1000 Genomes phase 1 unrelated data (1,083 individuals) [23]. As described earlier, PCs were first estimated in founders only, after LD pruning (including removing SNPs in long range high LD regions [24]) and merging with the 1000 Genomes dataset [23]. The non-founders were then projected into those PCs (eMethods5). Ancestry outliers were then removed based on visual inspection, using pairwise plots for the first seven principal components.

Similarly, PCA was used to identify substructure within each subpopulation of all MoBa batches. PCs were first estimated in founders only, after LD pruning (including removing SNPs in long range high LD regions [24]). The non-founders were projected into the PC space of the founders. Outliers were then removed based on visual inspection, using pairwise plots for the first ten principal components.

Finally, plate effect checks were performed, whereby: (1) the variation in PC1, PC2, PC3, and PC4 was investigated by creating box and whisker plots grouped by genotyping plate; (2) scatter plots were created for PC1 vs PC2 and PC3 vs PC4 coloured by genotyping plate to ensure clustering of plates did not occur; (3) Analysis of Variance (ANOVA) was performed to determine if there was a difference between genotyping plates within the first 10 PCs; and (4) the Cochran-Mantel-Haenszel test, in founders only, was used to assess the association of SNPs with genotyping plates (using the PLINK mh2 function with sex as the phenotype). SNPs associated with genotyping plates at a p-value < 0.0001 were removed.

#### Autosomal chromosomes

The QC was restricted to autosomal chromosomes at the beginning of the third round of the subpopulation QC (module 2). Therefore, a sex check was not performed in any of the remainder of module 2 or the remaining modules.

#### Sex chromosomes

QC, phasing, imputation, and post-imputation QC of chromosome X and pseudoautosomal regions (PAR) SNPs was performed separately from the autosomes. First, individuals were restricted to those passing module 9 post-imputation QC of the autosomal chromosomes. Then, the sex chromosome data was separated by chromosome (X, Y, and PAR) and by sex reported at birth for chromosome X and Y.

Basic QC was performed where SNPs and individuals were removed based on the following filters: (1) SNPs with MAF < 0.5%; (2) SNPs with call rate < 95%; (3) SNPs with call rate < 98% (eMethods2); (4) SNPs and individuals with a call rate < 98%; and (5) chromosome X SNPs in females only and PAR SNPs out of HWE at p < 1.00 × 10^−6^. A heterozygosity check was performed in PAR SNPs only. As heterozygosity was already checked in the autosomal data, individuals were only removed if the distribution is non-normal, given the small size of PAR.

Sex checks were then performed by first limiting to chromosome X and Y SNPs passing basic QC in both males and females, plus the PAR SNPs. Then a sex check was performed using (1) the chromosome X only and (2) chromosome Y only. First, the sex assignments reported at birth were checked against those imputed from X chromosome genotype calls and individuals with discordant sex were removed (females F > 0.2 and males < 0.8). Then the biological sex assignments were checked against the number of nonmissing Y chromosome genotype calls.

A ME check was then performed in families with one or two parents present in the data. Families with > 5% errors and SNPs with > 1% errors were removed. The remaining ME were set to missing.

Finally, a Cochran-Mantel-Haenszel test, in founders only, was used to assess the association of SNPs with genotyping plates (using the PLINK mh2 function with shuffled biological sex as the phenotype). SNPs associated with genotyping plates at a p-value < 0.0001 were removed.

### Module 3: Merge by genotyping array

Genotyping batches were merged by genotyping array (or genotyping arrays with significant overlap). Only SNPs passing three rounds of subpopulation QC in all the batches genotyped on the same array were kept. As multiple families were merged in each of the genotyping batches, the pedigree was updated to ensure the across-batch reported relationships were accurately coded in the merged datasets.

#### Sex chromosomes

Genotyping batches were merged by genotyping array (or genotyping arrays with significant overlap). Only SNPs passing the subpopulation QC in all batches genotyped on the same array were kept.

### Module 4: Subpopulation QC of merged genotyping array datasets

The subpopulation QC was then carried out in each merged genotyping dataset. The duplicate QC was performed as previously described, except both individuals from a true across-batch duplicate pair were kept, allowing this check to be performed in the post-imputation QC. All PCA were run using FlashPCA2.0 [26] to accommodate the increased number of individuals, with relatedness addressed as previously described. Whereby, PCs were first estimated in founders only, after LD pruning (including removing SNPs in long range high LD regions [14]). The non-founders were then projected into the PC space of the founders. The plate effect check was replaced with a batch effect check, whereby: (1) the variation in PC1, PC2, PC3, and PC4 was investigated by creating box and whisker plots grouped by genotyping batch; (2) scatter plots were created for PC1 vs PC2 and PC3 vs PC4 coloured by genotyping batch to ensure clustering of batches did not occur; (3) ANOVA was performed to determine if there was a difference in the first 10 PCs between genotyping batches; and (4) the Cochran-Mantel-Haenszel test, in founders only, was used to assess the association of SNPs with genotyping batches. SNPs associated with genotyping batches at a p-value < 0.0001 were removed (eMethods6).

#### Sex chromosomes

The subpopulation QC was then carried out in each merged genotyping dataset as described in module 2. The plate effect check was replaced with a batch effect check, whereby the Cochran-Mantel-Haenszel test, in founders only, was used to assess the association of SNPs with genotyping batches. SNPs associated with genotyping batches at a p-value < 0.0001 were removed.

### Module 5: Pre-phasing QC

A pre-phasing check was performed using the publicly available European Genome-Phenome Archive (Study ID EGAS00001001710) Haplotype Reference Consortium (HRC) release 1.1 [27] Imputation preparation and checking script version 4.2.13 [28], whereby, SNPs were removed if they were: (1) duplicate variants; (2) indels; (3) strand ambiguous (A/T and C/G); (4) not present among HRC sites; (5) > 0.2 minor allele frequency differences between the merged array MoBa dataset and HRC; and (6) alleles different to HRC. To match the data present in HRC, where needed, the following additional steps were implemented: (1) SNP IDs were updated to match the SNP IDs in HRC; (2) SNP alleles were flipped to match the strand of HRC; and (3) Allele 1 (A1) / allele 2 (A2) assignments were changed to match the A1 and A2 assignment in HRC.

### Module 6 & 7: Phasing and imputation

The publicly available European Genome-Phenome Archive (Study ID EGAS00001001710) HRC release 1.1 [27] data was used as the reference panel for both phasing and imputation. Whole-chromosome phasing [29] was performed using SHAPEIT2 release 904 [30] software with the duoHMM [31] algorithm to incorporate the pedigree information into the haplotype estimates. Imputation was performed using IMPUTE4.1.2_r300.3 [32] (eMethods7), an implementation of IMPUTE2 [33] re-coded to run faster with more memory efficiency, in chunks of up to 5,000 individuals (with all individuals from the same family kept in the same chunk). Imputation was conducted in five megabases chunks, with one megabase buffer on each side (meaning three megabases from each chunk were used for analyses). Chunks were defined from the first SNP passing pre-phasing QC on each chromosome. Imputation quality (INFO) scores were calculated by QCTOOL version 2.08 [34]. Initial, post-imputation checks were performed following the steps outlined in the ic tool version 1.0.8 [35].

#### X chromosome

The PAR (PAR1 and PAR2) were phased and imputed following the same protocols as those used for the autosomes. For the non-pseudo-autosomal region of the X chromosome, phasing was performed without application of the duoHMM algorithm, and imputation was carried out using IMPUTE2.3.2. INFO scores for this region were calculated using QCTOOL version 2.2.0. The rest of the processes mirrored the procedures applied to the autosomes.

### Module 8: Post-imputation QC

Post-imputation QC was performed in a single round following the QC by genotype array protocol for each imputation batch. Initially, dosage data was converted to best-guess, hard-call genotype data using a certainty threshold 0.7 (eMethods8). SNP and individual QC were then performed. The following thresholds were used for SNP removal: (1) imputation quality (INFO) score ≤ 0.8; (2) MAF < 1%; (3) call rate < 95% (eMethods9); (4) HWE p-value < 1.00 × 10^−6^; (5) discordant (concordance rate < 97%) in true duplicates (expected duplicates confirmed by genetic data PI_HAT > 0.95); (6) > 1% ME (with remaining ME set to missing); and (7) association with genotype batch at a p-value < 5 × 10^−8^. The following thresholds were used for individual removal: (1) call rate < 98%; (2) ± 3 standard deviations from the mean heterozygosity across all individuals; (3) the individual from each true duplicate pair with the lowest call rate (eMethods10); (4) all relatedness checks described in module 2; (5) cryptic relatedness; (6) ME > 5% in families; (7) ancestry outliers (PCA MoBa & 1000 Genomes [23]); and (8) subpopulation outliers (PCA MoBa). For the KING [25] relatedness checks temporary SNP removal was required due to computational resource limitations: SNPs with call rate < 98%, MAF < 5%, and pairs of SNPs with squared correlation greater than 0.4 were temporarily removed (eMethods11).

#### X chromosome

Post-imputation QC was performed in a single round following the QC by genotype array protocol for each imputation batch. Initially, the X and PAR dosage data was converted to best-guess, hard-call genotype data using a certainty threshold 0.7. The chromosome X data was then separated by sex reported at birth. Then the following thresholds were used for SNP removal: (1) SNPs with MAF < 1%; (2) call rate < 95%; (3) HWE at p < 1.00 × 10^−6^ in PAR and females only for chromosome X; (4) > 1% ME (with remaining ME set to missing); and (5) association with imputation batch at a p-value < 5 × 10^−8^. The following thresholds were used for individual removal: (1) call rate < 98%; (2) non-normal heterozygosity distribution of PAR; (3) ME > 5% in families; and (4) discordant sex (females F > 0.5 and males < 0.5).

### Module 9: Post-imputation QC of merged imputation batches

Post-imputation QC of the merged imputation batches is required for cohorts with multiple imputation batches to confirm all expected across batch relationships are as expected and ensure inferred across-imputation relationships are accurately coded in extended families (eMethods12). The imputation batches were merged keeping only overlapping SNPs passing post-imputation QC in all imputation batches. The pedigree was updated to ensure the across-imputation reported relationships were accurately coded in the merged dataset. The post-imputation SNP and individual QC procedures, including relatedness checks of updated across-imputation batch relationships, were then performed as described in module 8. The batch effect check was replaced with an imputation effect check, whereby: (1) the variation in PC1, PC2, PC3, and PC4 was investigated by creating box and whisker plots grouped by imputation batch; (2) scatter plots were created for PC1 vs PC2 and PC3 vs PC4 coloured by imputation batch to ensure clustering of imputation batches did not occur; (3) Analysis of Variance (ANOVA) was performed to determine if there was a difference in the first 10 PCs between imputation batches; and (4) the Cochran-Mantel-Haenszel test, in founders only, was used to assess the association of SNPs with imputation batches. SNPs associated with imputation batches at a p-value < 5 × 10^−8^ were removed.

#### X chromosome

The imputation batches were merged keeping only overlapping SNPs passing post-imputation QC in all imputation batches. The post-imputation SNP and individual QC procedures were then performed as described in module 8. The batch effect check was replaced with an imputation effect check, whereby the Cochran-Mantel-Haenszel test, in founders only, was used to assess the association of SNPs with imputation batches. SNPs associated with imputation batches at a p-value < 5 × 10^−8^ were removed.

## Results

In total, 238,001 MoBa samples were sent for genotyping in 26 batches. The 26 MoBa genotyping batches comprised 235,412 successfully genotyped samples. Of the successfully genotyped samples, 234,505 came from individuals still included in the study (i.e., who had not actively withdrawn from participating in MoBa) as of January 2019.

Approximately 95% of the participants were identified as having European ancestry based on PCA with 1000 Genomes (Module 1). Figure 2 shows PC clustering of individuals passing the MoBaPsychGen pipeline with the 1000 Genomes anchor (European, Asian, and African) populations.

**Figure 2.**
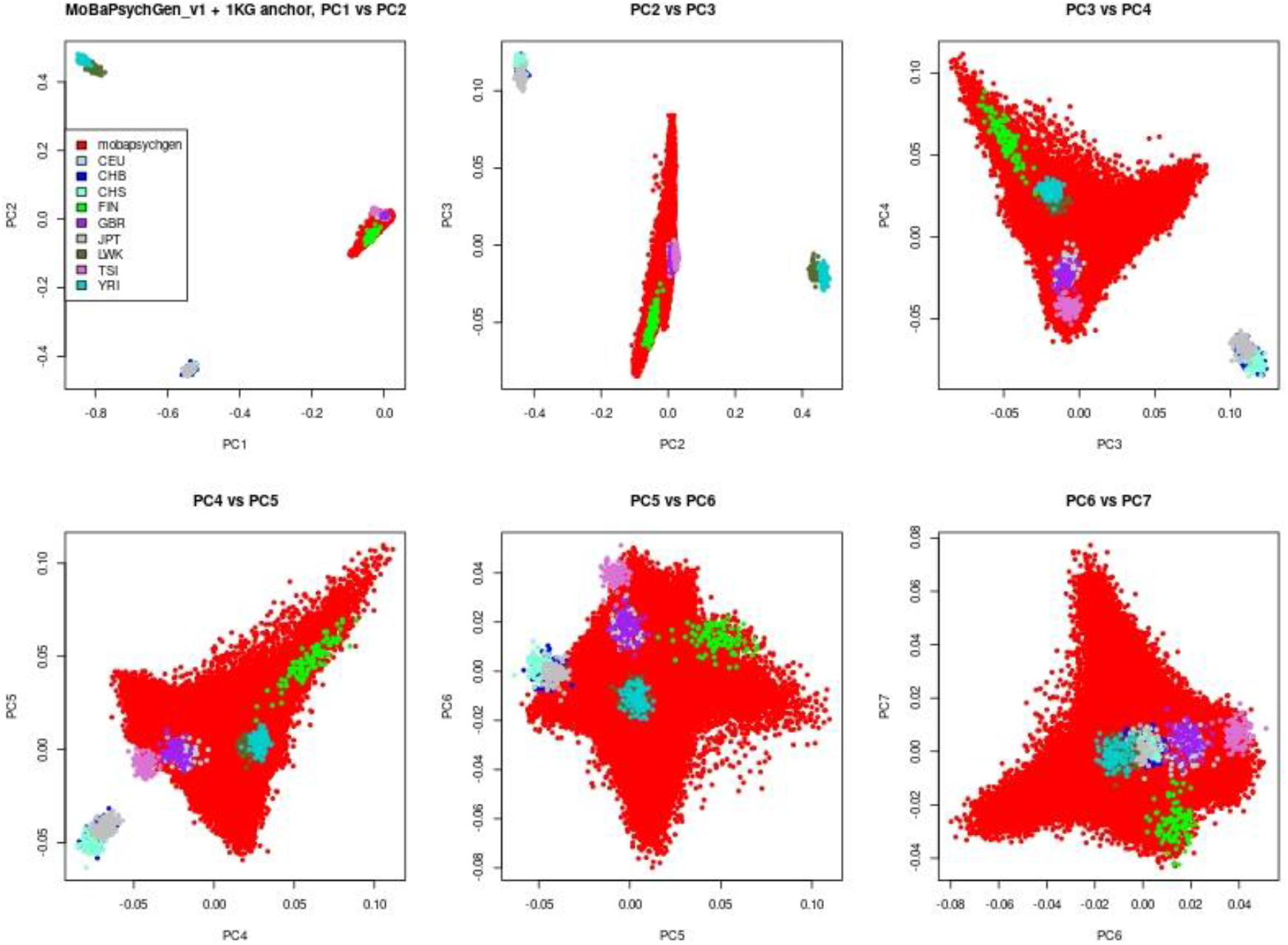
Plot of first 7 PCs of individuals passing MoBaPsychGen pipeline (red) with 1000 Genomes anchor (European, Asian, and African) populations. Legend abbreviations are: CEU = Utah residents with Northern and Western European ancestry, CHB = Han Chinese in Beijing, China, CHS = Southern Han Chinese, China, FIN = Finnish in Finland, GBR = British in England and Scotland, JPT = Japanese in Tokyo, Japan, LWK = Luhya in Webuye, Kenya, TSI = Toscani in Italy, and YRI = Yoruba in Ibadan, Nigeria.

After post-imputation QC the MoBaPsychGen pipeline output included 6,981,748 autosomal SNPs and 207,569 unique individuals (90% of the unique individuals included in the study), comprising 76,577 children, 53,358 fathers, and 77,634 mothers. A further 174,462 chromosome X and 3,200 PAR SNPs are available in a subset of these individuals (N = 204,913 and 135,593, respectively). An overview of the number of SNPs and individuals passing the various QC modules is available in Supplementary Tables 2-3 and imputation quality plots are available in Supplementary Figures 3-8.

Initially, the 234,505 samples (230,436 unique individuals) were assembled into 90,861 nuclear and blended families constructed based on the reported PO relationships for each pregnancy. The initial nuclear and blended family size based on reported PO relationships only ranged from singletons (where the other family members were not genotyped due to missing blood samples, genotyping failures, ect.) to eight unique individuals. The initial nuclear and blended families only included across generation PO relationships for each pregnancy, which enabled the identification of FS or HS in the child generation based on having the same reported parent (ie., does not include relationships inferred using genetic data).

The relatedness checks performed throughout the pipeline using the genetic data allowed identification (inferred from genetic data) of within-generation and across-generation first-degree (MZ, DZ, FS, PO), second-degree (HS, AUNN, GO), and third-degree (first cousins) relationships (Figure 3).

**Figure 3.**
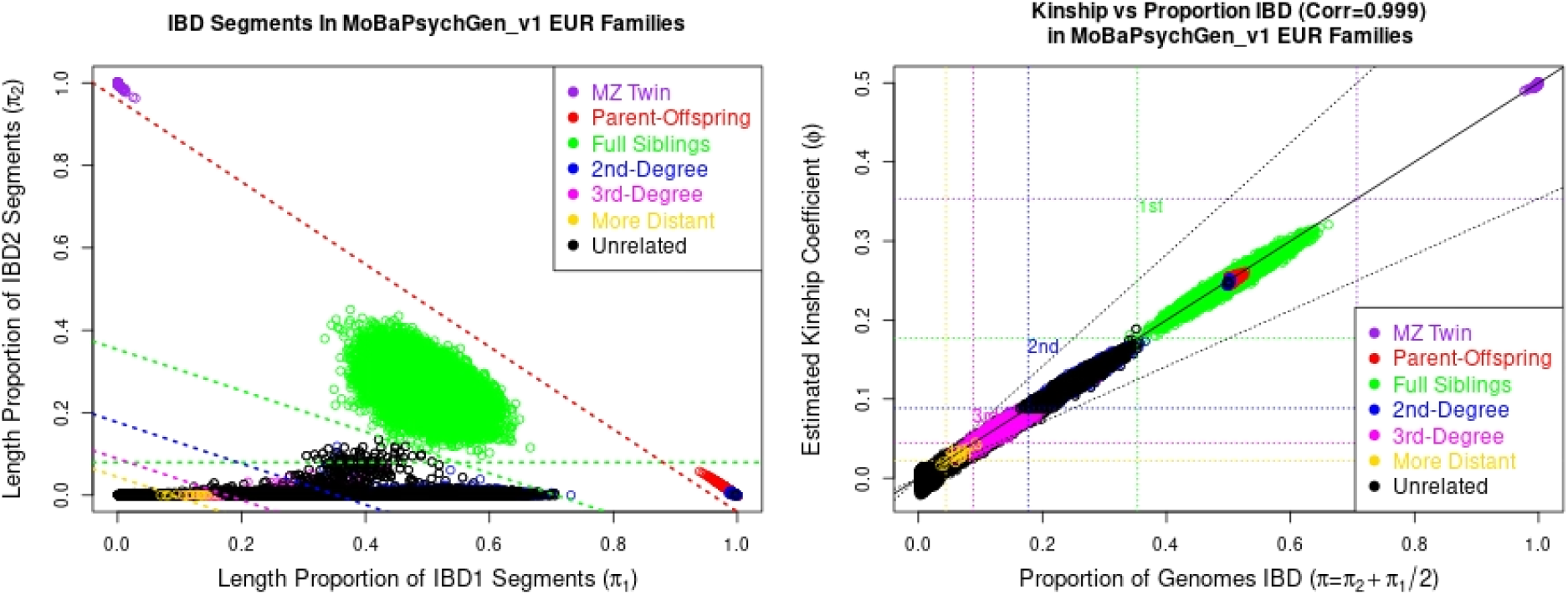
IBD plots showing the within-(extended) family relationships for the individuals passing MoBaPsychGen pipeline. The left plot shows the proportion of SNPs with 1 allele IBD vs the proportion of SNPs with 2 alleles IBD. The right plot shows the estimated proportion of the genome shared IBD vs the estimated kinship coefficient.

The individuals passing post-imputation QC comprised 64,471 nuclear, blended, and extended families ranging in size from singletons to 84 unique individuals (singletons are included as families as other family members may not have been genotyped, imputed, or passed post-imputation QC). The complexity of the pedigrees in MoBa is illustrated by the varying family sizes identified (Figure 4). The relationships identified included 287 monozygotic twin pairs, 22,884 full siblings, 117,004 PO pairs, 23,299 second-degree relative pairs, and 10,828 third-degree relative pairs. This included 44,017 full father-mother-child trios (with one complete trio in the parent generation), 4,592 father-child duos, and 24,380 mother-child duos.

**Figure 4.**
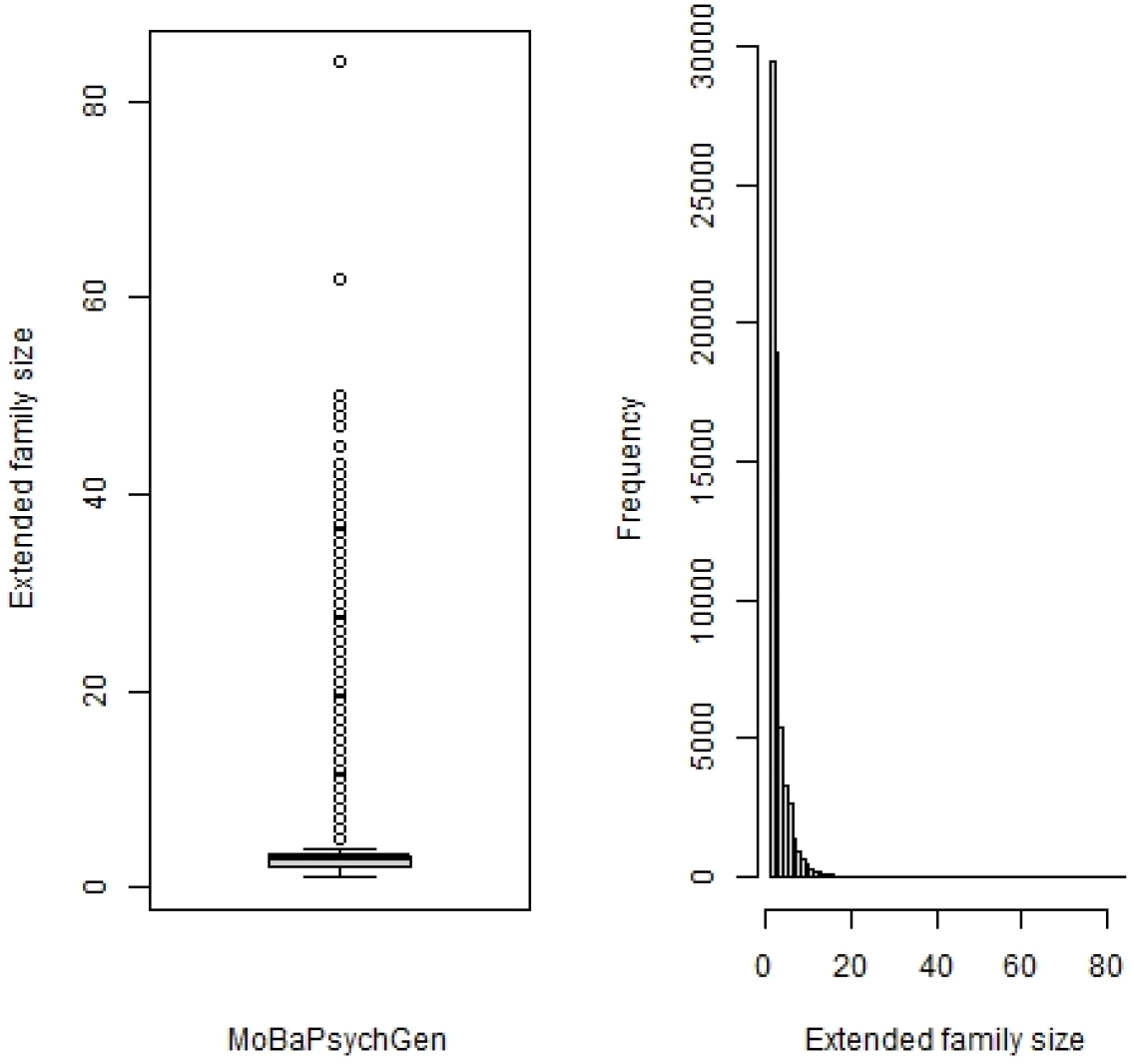
Plots showing the family size range of individuals passing MoBaPsychGen pipeline. Left: Box and whisker plot. Right: Histogram. Note, singletons are included in the family size range as other family members may not have been genotyped, imputed, or passed post-imputation QC.

## Discussion

We developed and applied a new QC and imputation pipeline capable of handling genotyping data from large population-based samples of individuals with complex interrelatedness. The MoBaPsychGen pipeline was designed for, and applied to MoBa, a cohort with a highly complex pattern of relatedness – over 84,500 first degree relatives, and approximately 12,000 second-degree relatives – and for which genotyping has been conducted across multiple research projects spanning several years, in batches selected based on varying criteria, and processed using different genotyping arrays at different genotyping centres. Individuals passing QC with the MoBaPsychGen pipeline represented 90% of the unique originally genotyped samples, with information on nearly 7 million SNPs available for all individuals in a final merged dataset.

Most currently available QC pipelines are structured to handle genotype data from unrelated individuals only or samples with a simple relatedness structure (e.g., nuclear families). As genotyping continues to decrease in cost and processing time, increasing numbers of people will be genotyped for research – including both in existing cohort samples with complex interrelatedness by design, and in convenience samples where complex relationships become inevitable as sample size increases (e.g., communities or small countries). Instead of excluding these valuable relationships, they can be leveraged in analyses to provide not only larger sample sizes, but also important discoveries, provided the data is processed in such a way that complex interrelatedness is accounted for and does not have the potential to bias results.

In developing the MoBaPsychGen pipeline, we focused on ensuring comprehensive coverage of key QC steps with particular importance for the MoBa sample. Firstly, the duplicate QC, which provides an indication of the heterogeneity within and between both genotyping batches and imputation batches. Through the removal of discordant SNPs, based on the relatively high number of duplicate or triplicate genotype samples in MoBa, we ensure that genotyping errors are identified and addressed. Secondly, the relatedness check, which corrects inaccurate pedigree assignment. With a high number of known relationships in cohort samples such as MoBa, it is possible to identify any sample mix-ups or unexpected duplicates – but doing so requires specific, manually-overseen steps in the QC pipeline. Furthermore, having as many relationships as possible coded in the pedigree is also important because: (1) only founders are used in some of the basic SNP QC steps (e.g., MAF and HWE) and (2) without correct pedigree information, the estimated haplotypes and resulting imputed genetic data will be affected, resulting in higher likelihood of switch errors in the haplotype estimation and ME in the imputed data [31]. It is particularly important that these key steps are performed accurately throughout the QC pipeline to ensure the highest quality data.

The MoBaPsychGen pipeline can be applied to other cohorts containing related individuals. Conducting quality control procedures on a cohort of genetically related individuals comes with inherent challenges. Added to this, additional complexities including duplicate samples, distinct genotyping batches of varying sizes based on varying selection criteria and processed on different genotyping arrays are idiosyncratic characteristics of the MoBa sample that present challenges for the QC process but which, nonetheless, may feature to varying degrees in other cohorts. The steps taken to meet these challenges in the MoBaPsychGen pipeline should provide an informative basis for those carrying out quality control in other large-scale population-based cohorts. Therefore, as individuals are recruited into cohorts in larger numbers, the prevalence of relatedness between participants and other complexities, such as having multiple genotyping batches, will increase. The MoBaPsychGen pipeline provides an example of how to handle these complexities.

The challenges and intricacies of using genomic data from samples with complex interrelatedness do not end with QC. Indeed, it is important that users understand the implications and decisions of the QC process and interact correctly with its outputs according to their analytic needs. To this end, for the MoBa genetic data to which the MoBaPsychGen pipeline was applied in this paper, and which is available in post-imputation QC form to MoBa researchers on request (see https://www.fhi.no/en/more/research-centres/psychgen/access-to-genetic-data-after-quality-control-by-the-mobapsychgen-pipeline-v/), we have developed an R package, *genotools* (https://github.com/psychgen/genotools) as a companion to the QCed genetic data. This package aims to facilitate accurate, efficient, and reproducible use of MoBa genetic data, and currently has functionality to help users work with genetic datafiles alongside supporting identification, linkage, and covariates files, identifying relationships based on analysis types, creating and adjusting polygenic scores, and will continue to be developed to help researchers maximize the potential of analyses using MoBa genetic data processed via the MoBaPsychGen pipeline.

Many novel methods can make use of genetic data from related individuals to gain additional insights into the nature of genetic effects that are not accessible using population-based data. These include variations of GWA analyses (e.g., within-family GWA, sibling GWA, and trio GWA analyses), genomic-relatedness matrix-based approaches (e.g., Trio GCTA, M-GCTA [36]), polygenic score-based approaches (e.g., trio polygenic score analyses, transmitted and non-transmitted allele polygenic score analyses, rGenSi modelling [37], polygenic transmission disequilibrium testing) and others (e.g., within-family Mendelian Randomization). It is likely that methodological development in this space will continue at a high rate over the coming years. Appropriately preparing genomic data in relevant samples for inclusion in analyses using this wide array of approaches is an essential upstream component of research into a range of important topics, including familial aggregation of risk for health problems, putative causal risk factors (including parental factors) in children’s environments, comorbidity between health problems, factors influencing early life neurodevelopment and many more. Furthermore, knowledge about the specific degrees of relatedness within a cohort allows for contribution to larger consortia of certain family types, for example, sibling, twin, and children of twin consortia.

## Limitations and future directions

Every effort was taken to ensure the MoBaPsychGen pipeline adhered to the current best-practice protocols for preforming QC while appropriately handling the idiosyncratic characteristics of the MoBa genotyping data. However, there are a few limitations of the pipeline. Namely, that the pipeline has only been implemented for autosomes in the largest subpopulation of MoBa, which is the European subpopulation. We are currently working on updating the pipeline to address these limitations by completing the QC and imputation of individuals from the other core subpopulations.

## Conclusion

The MoBaPsychGen pipeline handles complex relatedness and other specific challenges in the process of quality controlling genotype data from the MoBa cohort. The pipeline has potential utility in similar cohorts, and MoBa genetic data processed via the pipeline is available and well-suited to a wide range of highly informative downstream genetic analyses - including those that make particular use of the complex relatedness in such a sample.

## Supporting information

MoBaPsychGen_suppl

## Acknowledgements

We are grateful to all the participating families in Norway who take part in this on-going cohort study. Many thanks to Yunhan Chu for her analytic and scripting contribution to the QC. This work was performed on the Tjeneste for Sensitive Data (TSD) facilities, owned by the University of Oslo, operated and developed by the TSD service group at the University of Oslo, IT-Department (USIT), using resources provided by Sigma2—the National Infrastructure for High Performance Computing and Data Storage in Norway (UNINETT). Thank you to Raymond Walters and Stephan Ripke for their advice on the QC.

## Code and data availability

Scripts used throughout the MoBaPsychGen pipeline are available on GitHub: https://github.com/psychgen/MoBaPsychGen-QC-pipeline. Information on how to access the MoBaPsychGen post-imputation QC data is available here: https://www.fhi.no/en/more/research-centres/psychgen/access-to-genetic-data-after-quality-control-by-the-mobapsychgen-pipeline-v/.

