## Supplementary material for "The Norwegian Mother, Father, and Child cohort study (MoBa) genotyping data resource: MoBaPsychGen pipeline v.1": MoBaPsychGen_suppl

### eMethods1 – genotyping of MoBa

#### Genotyping in the HARVEST and SELECTIONpreDISPOSED projects

Two HARVEST batches, comprising approximately 11,000 full mother, father, and child trios (approximately 33,000 individuals), were genotyped in 2014 and 2015. Genotyping, following the Illumina Infinium HD ultra-protocol with scanning performed using the Illumina iScan, and the conversion of scanned data to genotype calls, using GenomeStudio_v2011.1 with GenTrain_v2, was conducted at the Genomics Core Facility, Trondheim, Norway. Two Illumina HumanCoreExome arrays were used to genotype the batches. The HARVEST12 batch was genotyped using the Illumina HumanCoreExome12v1.1 array. The HARVEST24 batch was genotyped using the Illumina HumanCoreExome24v1.0 array. Because of a difference in signal intensities, the HARVEST12 batch was re-clustered into two sub-batches: HARVEST12a and HARVEST12b. Additionally, the HapMap sample NA12878 was genotyped twice as an external control.

An additional 6,000 full trios were genotyped in 2017 (ROTTERDAM1 batch) and a further 3,000 trios were genotyped in 2018 (ROTTERDAM2 batch). Collectively, these two batches comprised approximately 9,000 full mother, father, and child trios (approximately 27,000 individuals). Genotyping using the Illumina Global Screening Array MD v.1.0 array and the conversion of scanned data to genotype calls using GenomeStudio was conducted at the Erasmus MC, Rotterdam, Netherlands.

The full trios were selected randomly, while minimizing the number of trays taken out of the freezer, but were excluded if: (1) the pregnancy resulted in a stillbirth; (2) a sample from the trio was deceased (at the time the sample list was created); (3) the pregnancy resulted in a multiple birth (multiple births in the parent generation were not excluded); (4) data for the pregnancy is missing in the MBRN data; (5) missing anthropometric measurements at birth in the MBRN; (6) not included in the mother-reported first questionnaire (Q1) (as a proxy for higher fallout rate); and (7) missing DNA samples.

#### Genotyping in the NORMENT project

Genotyping of 20 batches with varying selection criteria and genotype arrays, and GenomeStudio conversion of scanned data to genotype calls were conducted at deCODE Genetics in Reykjavik, Iceland.

##### ADHD batches

Two data selections with the same criteria were conducted for the Common and rare genetic risk factors for ADHD subproject. The initial selection criteria were: (1) data from the MBRN available for the index pregnancy (the pregnancy with an ADHD diagnosis); (2) DNA available for the index pregnancy; (3). index pregnancy not stillborn or deceased; and (4) DNA available for the parents. Additional criteria for: (1) case children–ADHD diagnosis (International Classification of Diseases version 10 code F90) from Norwegian Patient Registry (NPR) which contains all diagnosis from specialist health service in Norway from 2008 onwards; (2) control children–no ADHD diagnosis in NPR plus birth year, sex, and birth hospital or county matched with case children; (3) mothers of case children–DNA available from mother at any time point, including from another pregnancy; and (4) fathers of case children–DNA available from father at any time point, might come from another pregnancy, and independent of DNA availability from the mother. The ADHD1 batch included 5,818 individuals, which were sent for genotyping using the Illumina InfiniumOmniExpress-24v1.2 array. The ADHD2 batch included 2,502 individuals, which were sent for genotyping using the Illumina GlobalScreening array MD v.1.0 array.

##### NORMENT parent only batches

The NORMENT-Jan2015 and NORMENT-Jun2015 batches comprised 6,040 unrelated parent samples, which were sent to be genotyped as controls for planned analyses, using the Illumina HumanOmniExpress-24v1.0 array.

##### NORMENT trio batches

The NORMENT-May16 batch included approximately 6,452 full trios, which were sent for genotyping using the Illumina InfiniumOmniExpress-24v1.2 array. Similarly, NORMENT-Feb18 batch included approximately 3,280 full trios, which were sent for genotyping using the Illumina Global Screening Array MD v.1.0 array.

##### NORMENT remaining MoBa samples

From 2020 onwards, the genotyping batches contained as many individuals from the same families as possible. However, the primary goal was to efficiently genotype the remaining samples from the MoBa cohort, with no selection criteria enforced other than having already been genotyped (meaning stillborn and deceased pregnancies or deceased individuals will be present in these batches). Therefore, the percentage of relatedness within the batches decreased with time.

The NORMENT-Feb2020 batch included 17,930 individuals with a moderate percentage of first-degree relatives, which was sent for genotyping using the Illumina Global Screening Array. Due to the availability of availability the NORMENT-Feb2020 batch was genotyped using two versions of the array and was therefore separated into two sub-batches: NORMENT-Feb2020_v1 and NORMENT-Feb2020_v3. The majority (13,505) of samples were genotyped using the Illumina Global Screening Array MD v.1.0, forming the NORMENT-Feb20-v1 sub-batch. However, a quarter of the samples (4,418) were genotyped using the Illumina Global Screening Array MD v.3.0, forming the NORMENT-Feb20-v3 sub-batch.

All remaining batches were genotyped using the Illumina Global Screening Array MD v.3.0 array and contained only a small percentage of first-degree relatives, with the last batch containing unrelated individuals only. The remaining 13 batches, NORMENT-Aug2020_996, NORMENT-Aug2020_1029, NORMENT-Nov2020_1066, NORMENT-Nov2020_1077, NORMENT-Nov2020_1108, NORMENT-Nov2020_1109, NORMENT-Nov2020_1135, NORMENT-Nov2020_1146, NORMENT-Mar2021_1273, NORMENT-Mar2021_1409, NORMENT-Mar2021_1413, NORMENT-Mar2021_1531, NORMENT-Mar2021_1532, comprised 25,004, 25,004, 25,002, 4,700 4,794, 5,640, 4,606, 5,264, 5,450, 2,703, 5,639, 1,974, and 219 samples sent for genotyping, respectively.

### eMethods2 – call rate filtering

When PLNK is run including a call rate filter for both SNPs (--geno) and individuals (--mind) in the same command the individuals filtering is performed first. To try and keep as many samples as possible the SNP call rate filtering with a threshold of 98% was performed twice, firstly in a PLINK command alone to remove SNPs with low call rate, and secondly, in a PLINK command with the individual call rate filter; with the second command initially removing and individuals with a low call rate before ensuring no SNPs have a low call rate after individuals with high missingness have been removed.

### eMethods3 – heterozygosity outlier filtering

Heterozygosity filtering was performed using a threshold of ± 3 standard deviations from the mean heterozygosity across all individuals rather than an F statistic to ensure the hetrozogosity distribution was normally distributed in all genotyping batches. In large genotyping batches the F statistic plots were normally distributed, however, in small genotyping batches large tails were observed (Supplementary Figure 1), however, limited or no outliers were removed. Therefore, filtering using ± 3 standard deviations from the mean heterozygosity across all individuals was used to ensure heterozygosity outliers were removed.

Supplementary Figure 1. Heterozygosity plots from the ADHD1 batch. Left plot shows the heterozygosity rate vs number of missing SNPs, with red mean ± 3 SD thresholds. Middle plot shows shows the Fhet vs Frequency plot. Right plot shows the Fhet vs Frequency plot with red Fhet | 0.2 | thresholds.

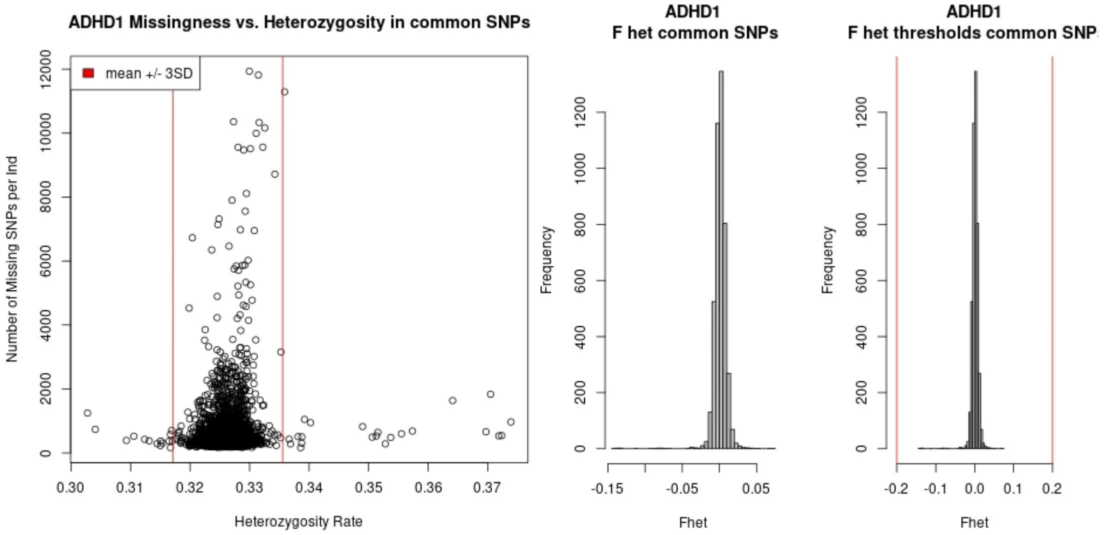

### eMethods4 –pedigree build and relatedness check software

KING was used to perform the pedigree build and relatedness checks instead or PLINK because (1) In the presence of admixture KING can accurately infer first-degree, second-degree, and third-degree relationships; and (2) preforms a pedigree build and outputs files to easily update family and parental IDs, whereas using PLINK the pedigree build would need to be manually curated likely and would likely introduce errors in such a large cohort.

### eMethods5 – PCA with 1000 Genomes and MoBa

Comparison of PCs estimated in 1000 Genomes with MoBa projected into those PCs and PCs estimated in 1000 Genomes plus MoBa founders with MoBa non-founders shows the first 5 PCs are highly correlated. In the NORMENT_Nov2020_1146 batch the correlations for PC1, PC2, PC3, PC4, and PC5 were 0.9998, -0.9994, 0.9944, 0.9854, and 0.9301, respectively.

### eMethods6 – batch effect checks

The two ADHD batches contained child cases and their parents, plus matched control children. Furthermore, seven batches were genotyped as full (mother-father-child) trios. Therefore, a concern of the batch effect check is that some variants associated with ADHD or trio-ness could be removed via selection biases. However, the batch effect check was performed in founders only and the number of SNPs removed due to association with batch was small (0.10 - 0.22% of SNPs removed). Furthermore, looking at the PC1 vs PC2 and PC3 vs PC4 plots coloured by batch the SNP removal is driven by large variation in sample size rather than clustering of specific batches (Supplementary Figure 2).

Supplementary Figure 2. PC1 vs PC2 and PC3 vs PC4 plots coloured by batch (black: ADHD2, red: NORMENT_Mar2021_1273, dark blue: NORMENT_Mar2021_1409, green: NORMENT_Mar2021_1413, light blue: NORMENT_Mar2021_1531, pink: NORMENT_Mar2021_1532).

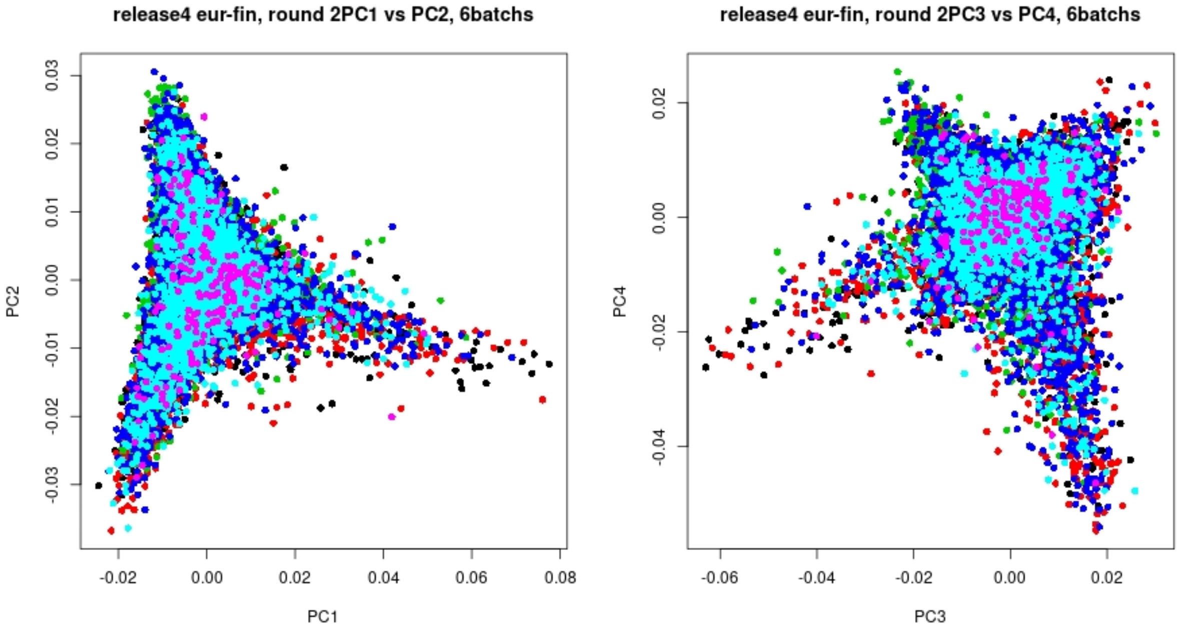

### eMethods7 – imputation software

IMPUTE4 was used for imputation as it uses the methodology implemented in IMPUTE2, which can handle both related and unrelated individuals, with more computational efficiency, as it was specifically designed for the imputation of large datasets. IMPUTE5 was not considered as we were unsure whether the software can appropriately account for varying levels of relatedness during imputation.

#### eMethods8 – genotype certainty threshold

The certainty threshold of 0.7 was used to balance the high missingness that results from using the PLINK default of 0.9 and with the level of uncertainty that is added to the dataset by lowering the threshold.

#### eMethods9 – post-imputation QC SNP call rate filter

The post-imputation QC SNP call rate filter was reduced to 95% as researchers may have specific reasons for investigating SNPs with slightly lower SNP call rate threshold. Therefore, we have applied a lower SNP call rate threshold of 95% during the post-imputation QC. If researchers require the higher SNP call rate threshold of 98%, they can easily apply this to the data using plink.

#### eMethods10 – post-imputation QC duplicate removal

Genotyping batch was not used as the as the deciding factor to identify which individual from each duplicate pair to remove, because the varying selection criteria of genotyping batches prevented all nuclear and blended family members being genotyped in the same genotyping batch. Meaning it is not possible to preserve the integrity of genotyping array within families. Therefore, we chose to use call rate as the deciding factor to identify the duplicate to remove to prevent introducing biases on which family members to have genotyped using the same array.

#### eMethods11 – filtering imputed SNPs for KING analyses

KING website (https://www.kingrelatedness.com/manual.shtml) recommends: "Please do not prune or filter any good SNPs that pass QC prior to any KING inference, unless the number of variants is too many to fit the computer memory, in which case rare variants can be filtered out. LD pruning is not recommended in KING". However, this is referring to directly genotyped SNPs. To get enough SNP filtering a really high MAF threshold had to be used (approx. > 30%). Therefore, given the SNPs were imputed light LD pruning was used to ensure the SNPs were distributed across the genome, rather than INFO score filtering, which is correlated with MAF, and would have led to an uneven distribution of SNPs across the genome (with SNPs in high LD with directly genotyped SNPs likely to have higher INFO scores).

#### eMethods12 – merge imputation batches

MoBa was genotyped through multiple projects with varying selection criteria. As a result, the reported nuclear and blended families were not genotyped as complete families. Therefore, to ensure whole families can be analysed together and MoBa can be analysed as a complete cohort, the imputation batches were merged. Merging only overlapping SNPs passing post-imputation QC in all imputation batches ensures a single distribution of power and non-centrality parameter across all SNPs. Furthermore, a central assumption of standard meta-analysis approaches is that effect sizes are independent. Methodologies are being developed to conduct meta-analyses of results with non-independent effect estimates, however, there are significant limitations to these methods [1].

#### Supplementary text References

1. Cheung, M.W.L., *A Guide to Conducting a Meta-Analysis with Non-Independent Effect Sizes.* Neuropsychology Review, 2019. **29**(4): p. 387-396.

Supplementary Table 1. Overview of the MoBa genotyping batches.

| Project | Genotyping batch | Number of samples sent for genotyping | Number of samples successfully genotyped | | Number of genotyped SNPs | Genotyping center | Genotyping array | Genotype build | Basic selection criteria |
|---|---|---|---|---|---|---|---|---|---|
| Novel Tools for Childhood Predisposition to Obesity and Diabetes (SELECTIONpreDISPOSED) | harvest12 | 20,668 (including 250 within-batch duplicates) | harvest12a sub-batch | 18,972 | 542,585 | Genomics Core Facility, Trondheim, Norway | Illumina HumanCoreExome12v1.1 | GRCh37 / hg19 | Full Trios |
| | | | harvest12b sub-batch | 1,692 | 542,585 | | | | |
| Better Health by Harvesting Biobanks (HARVEST) | harvest24 | 12,874 (including 138 within-batch duplicates) | 12,874 | | 547,644 | | Illumina HumanCoreExome24v1.0 | | |
| SELECTIONpreDISPOSED | rotterdam1 | Approximately 18,000 | 17,949 | | 692,367 | ERASMUS MC, Rotterdam, Netherlands | Illumina Global Screening Array MD v.1.0 | | |
| | rotterdam2 | Approximately 9,000 | 9,041 | | 692,338 | | | | |
| NORMENT sub-projects | norment-feb-2018 | 9,841 | 9,632 | | 693,143 | deCODE Genetics, Reykjavik, Iceland | | GRCh38 / hg38 | |
| | norment-jan-2015 | 6,040 | 2,983 | | 710,146 | | Illumina HumanOmniExpress-24v1.0 | GRCh37 / hg19 | Unrelated parents genotyped as controls |
| | norment-jun-2015 | | 2,976 | | 708,882 | | | | |
| | norment-may-2016 | 19,357 | 17,608 | | 712,628 | | Illumina InfiniumOmniExpress-24v1.2 | GRCh38 / hg38 | Full trios |
| | adhd1 | 5,790 | 5,410 | | 713,599 | | | GRCh37 / hg19 | ADHD child case trios and singleton controls |
| | adhd2 | 2,502 | 2,426 | | 693,143 | | Illumina Global Screening Array MD v.1.0 | GRCh38 / hg38 | |
| | norment-feb-2020 | 17,930 | norment-feb-2020_v1 sub-batch | 13,505 | 693,143 | | Illumina Global Screening Array MD v.1.0<br>Illumina Global Screening Array MD v.3.0 | | Unrelated individuals with moderate percentage of |

| | | | | | | | | | |
|---|---|---|---|---|---|---|---|---|---|
| | | | norment-feb-2020_v3 sub-batch | 4,418 | 687,31 | | | | first-degree relatives |
| | norment-aug-2020_996 | 25,004 | 25,499 | | 687,316 | | Illumina Global Screening Array MD v.3.0 | | Unrelated individuals with small percentage of first-degree relatives |
| | norment-aug-2020_1029 | 25,004 | 24,980 | | 687,316 | | | | |
| | norment-nov-2020_1066 | 25,002 | 24,995 | | 687,316 | | | | |
| | norment-nov-2020_1077 | 4,700 | 4,699 | | 687,316 | | | | |
| | norment-nov-2020_1108 | 4,794 | 4,792 | | 687,316 | | | | |
| | norment-nov-2020_1109 | 5,640 | 5,625 | | 687,316 | | | | |
| | norment-nov-2020_1135 | 4,606 | 4,605 | | 687,316 | | | | |
| | norment-nov-2020_1146 | 5,264 | 5,256 | | 687,316 | | | | |
| | norment-mar-2021_1273 | 5,450 | 5,446 | | 687,316 | | | | |
| | norment-mar-2021_1409 | 2,703 | 2,702 | | 687,316 | | | | |
| | norment-mar-2021_1413 | 5,639 | 5,637 | | 687,316 | | | | |
| | norment-mar-2021_1531 | 1,974 | 1,971 | | 687,316 | | | | |
| | norment-mar-2021_1532 | 219 | 219 | | 687,316 | | | | Unrelated |

Supplementary Table 2. Number of autosomal SNPs and Individuals passing various stages of the QC pipeline

| Imputation batch | Genotyping batch / sub-batch | Number of SNPs | | | | | Number of Individuals | | | | | | | |
|---|---|---|---|---|---|---|---|---|---|---|---|---|---|---|
| | | Genotyped | Pass QC (Module 2) | Pass merged QC (Module 4) | Pass post-imputation QC (Module 8) | Pass merged post-imputation QC (Module 9) | Sent for genotyping | Genotyped | European subpopulation selection (Module 1) | Pass QC (Module 2) | Pass merged QC (Module 4) | Imputed (Module 6 & 7) | Pass post-imputation QC (Module 8) | Pass merged post-imputation QC (Module 9) |
| HCE | harvest12b | 516,071 | 270,779 | 257,576 | 7,210,134 | 6,981,748 | 20,668 (250 within-batch duplicates) | 1,692 | 1,605 | 1,546 | 30,859 | 30,859 | 30,735 | 207,569 |
| | harvest12a | 516,602 | 273,345 | | | | | 18,972 | 18,079 | 17,509 | | | | |
| | harvest24 | 523,191 | 278,433 | | | | 12,874 (138 within-batch duplicates) | 12,874 | 12,409 | 12,002 | | | | |
| GSA1 | rotterdam1 | 665,793 | 495,425 | 456,904 | 7,476,429 | | Approx. 18,000 | 17,949 | 17,143 | 16,662 | 33,893 | 33,893 | 33,853 | |
| | rotterdam2 | 665,764 | 502,966 | | | | Approx. 9,000 | 9,041 | 8,689 | 8,403 | | | | |
| | norment-feb-2018 | 665,764 | 536,366 | | | | 9,841 | 9,632 | 9,319 | 9,078 | | | | |
| OMNI | norment-jan-2015 | 690,723 | 600,361 | 565,795 | 7,479,229 | | 6,040 | 2,983 | 2,886 | 2,812 | 26,891 | 26,891 | 26,501 | |
| | norment-jun-2015 | 689,506 | 591,946 | | | | | 2,976 | 2,763 | 2,685 | | | | |
| | norment-may-2016 | 693,173 | 612,994 | | | | 19,357 | 17,608 | 17,074 | 16,666 | | | | |
| | adhd1 | 693,521 | 613,042 | | | | 5,790 | 5,410 | 5,075 | 4,934 | | | | |
| GSA2 | norment-feb-2020_v1 | 673,639 | 538,857 | 481,783 | 7,503,545 | | 17,930 | 13,505 | 12,962 | 12,639 | 62,181 | 62,181 | 62,013 | |
| | norment-feb-2020_v3 | 654,773 | 538,857 | | | | | 4,418 | 4,192 | 4,069 | | | | |

| | | | | | | | | | | | | | |
|---|---|---|---|---|---|---|---|---|---|---|---|---|---|
| | norment-aug-2020_996 | 654,773 | 540,325 | | | | 25,004 | 24,999 | 23,850 | 23,227 | | | |
| | norment-aug-2020_1029 | 654,773 | 541,047 | | | | 25,004 | 24,980 | 23,654 | 23,115 | | | |
| GSA3 | norment-nov-2020_1066 | 654,773 | 531,050 | 512,910 | 7,513,030 | | 25,002 | 24,995 | 23,741 | 23,141 | 45,624 | 45,624 | 45,563 |
| | norment-nov-2020_1077 | 654,773 | 530,344 | | | | 4,700 | 4,699 | 4,462 | 4,330 | | | |
| | norment-nov-2020_1108 | 654,773 | 528,681 | | | | 4,794 | 4,792 | 4,543 | 4,465 | | | |
| | norment-nov-2020_1109 | 654,773 | 527,482 | | | | 5,640 | 5,625 | 5,324 | 5,199 | | | |
| | norment-nov-2020_1135 | 654,773 | 526,353 | | | | 4,606 | 4,605 | 4,344 | 4,265 | | | |
| | norment-nov-2020_1146 | 654,773 | 525,685 | | | | 5,264 | 5,256 | 4,942 | 4,808 | | | |
| GSA4 | norment-mar-2021_1273 | 654,773 | 513,638 | 459,765 | 7,484,274 | | 5,450 | 5,446 | 5,131 | 5,018 | 16,768 | 16,768 | 16,758 |
| | norment-mar-2021_1409 | 654,773 | 508,297 | | | | 2,703 | 2,702 | 2,595 | 2,507 | | | |
| | norment-mar-2021_1413 | 654,773 | 520,646 | | | | 5,639 | 5,637 | 5,330 | 5,184 | | | |
| | norment-mar-2021_1531 | 654,773 | 517,471 | | | | 1,974 | 1,971 | 1,853 | 1,800 | | | |

| | | | | | | | | | | |
|---|---|---|---|---|---|---|---|---|---|---|
| | norment-mar-2021_1532 | 654,773 | 494,207 | | | | 219 | 219 | 205 | 196 |
| | adhd2 | 673,639 | 534,528 | | | | 2,502 | 2,426 | 2,279 | 2,199 |
| NB the large reduction in the number of SNPs passing QC in the batches (harvest12a, harvest12b, and harvest24) is due to the coverage of the HumanCoreExome array, which has a very high proportion of rare SNPs. | | | | | | | | | | |

Supplementary Table 3. Number of chromosome X and PAR SNPs and individuals passing various stages of the QC pipeline

| Imputation batch | Genotyping batch / sub-batch | Number of SNPs (chromosome X / PAR) | | | | | Number of Individuals (chromosome X / PAR) | | | | | |
|---|---|---|---|---|---|---|---|---|---|---|---|---|
| | | Genotyped | Pass QC (Module 2) | Pass merged QC (Module 4) | Pass post-imputation QC (Module 8) | Pass merged post-imputation QC (Module 9) | Pass autosomal merged post-imputation QC (Module 9) | Pass QC (Module 2) | Pass merged QC (Module 4) | Imputed (Module 6 & 7) | Pass post-imputation QC (Module 8) | Pass merged post-imputation QC (Module 9) |
| HCE | harvest12b | 12,853 / 150 | 7,155 / 21 | 6,883 / 16 | 193,410 / NA | 174,462 / 3,200 | 1,327 | 1,283 | 26,421 | 26,421 / NA | 26,421 / NA | 204,913 / 135,593 |
| | harvest12a | 12,862 / 150 | 7,380 / 39 | | | | 15,159 | 14,844 | | | | |
| | harvest24 | 12,834 / 146 | 7,308 / 22 | | | | 10,535 | 10,295 | | | | |
| GSA1 | rotterdam1 | 17,790 / 576 | 10,469 / 407 | 10,034 / 386 | 198,540 / 3,590 | | 15,672 | 15,560 | 32,289 | 32,289 | 322,289 / 29,213 | |
| | rotterdam2 | 17,790 / 576 | 11,135 / 411 | | | | 8,152 | 8,094 | | | | |
| | norment-feb-2018 | 17,385 / 524 | 12,654 / 396 | | | | 8,711 | 8,638 | | | | |
| OMNI | norment-jan-2015 | 17,441 / 682 | 13,189 / 360 | 12,001 / 278 | 207,078 / NA | | 2,715 | 2,714 | 26,004 | 26,004 / NA | 26,004 / NA | |
| | norment-jun-2015 | 17,395 / 681 | 12,724 / 354 | | | | 2,611 | 2,570 | | | | |
| | norment-may-2016 | 17,718 / 480 | 13,823 / 314 | | | | 16,373 | 16,248 | | | | |
| | adhd1 | 17,565 / 683 | 14,037 / 366 | | | | 4,651 | 4,501 | | | | |
| GSA2 | norment-feb-2020_v1 | 17,385 / 524 | 12,859 / 396 | 10,746 / 369 | 201,987 / 3,389 | | 12,287 | 12,201 | 60,163 | 60,163 | 60,163 / 53,492 | |
| | norment-feb-2020_v3 | 27,064 / 729 | 20,589 / 594 | | | | 3,923 | 3,891 | | | | |
| | norment-aug-2020_996 | 27,064 / 729 | 19,913 / 586 | | | | 22,241 | 22,010 | | | | |

| | | | | | | | | | | | |
|---|---|---|---|---|---|---|---|---|---|---|---|
| | norment-aug-2020_1029 | 27,064 / 729 | 19,973 / 582 | | | | 22,296 | 22,103 | | | |
| GSA3 | norment-nov-2020_1066 | 27,064 / 729 | 20,610 / 592 | 18,909 / 571 | 217,230 / 5,263 | | 22,431 | 22,227 | 44,139 | 44,139 | 44,139 / 40,817 |
| | norment-nov-2020_1077 | 27,064 / 729 | 20,309 / 591 | | | | 4,165 | 4,099 | | | |
| | norment-nov-2020_1108 | 27,064 / 729 | 19,988 / 592 | | | | 4,294 | 4,207 | | | |
| | norment-nov-2020_1109 | 27,064 / 729 | 19,747 / 592 | | | | 5,000 | 4,926 | | | |
| | norment-nov-2020_1135 | 27,064 / 729 | 19,702 / 587 | | | | 4,162 | 4,094 | | | |
| | norment-nov-2020_1146 | 27,064 / 729 | 19,601 / 589 | | | | 4,661 | 4,593 | | | |
| GSA4 | norment-mar-2021_1273 | 27,064 / 729 | 18,906 / 583 | 8,699 / 365 | 197,640 / 3,483 | | 4,829 | 4,761 | 15,897 | 15,897 | 15,897 / 14,054 |
| | norment-mar-2021_1409 | 27,064 / 729 | 18,131 / 583 | | | | 2,409 | 2,388 | | | |
| | norment-mar-2021_1413 | 27,064 / 729 | 19,416 / 585 | | | | 5,005 | 4,946 | | | |
| | norment-mar-2021_1531 | 27,064 / 729 | 18,881 / 588 | | | | 1,741 | 1,704 | | | |
| | norment-mar-2021_1532 | 27,064 / 729 | 16,689 / 583 | | | | 182 | 179 | | | |
| | adhd2 | 17,385 / 524 | 12,514 / 393 | | | | 1,946 | 1,932 | | | |

NB the large reduction in the number of SNPs passing QC in the batches (harvest12a, harvest12b, and harvest24) is due to the coverage of the HumanCoreExome array, which has a very high proportion of rare SNPs.

Supplementary Figure 3. Imputation quality plots showing chromosome position vs imputation quality scores (INFO) for each chromosome and MoBa imputation batch. Plots from left to right are Imputation batch HCE, OMNI, GSA1, GSA2, GSA3, and GSA4. Plots from top to bottom are chromosome 1-22.

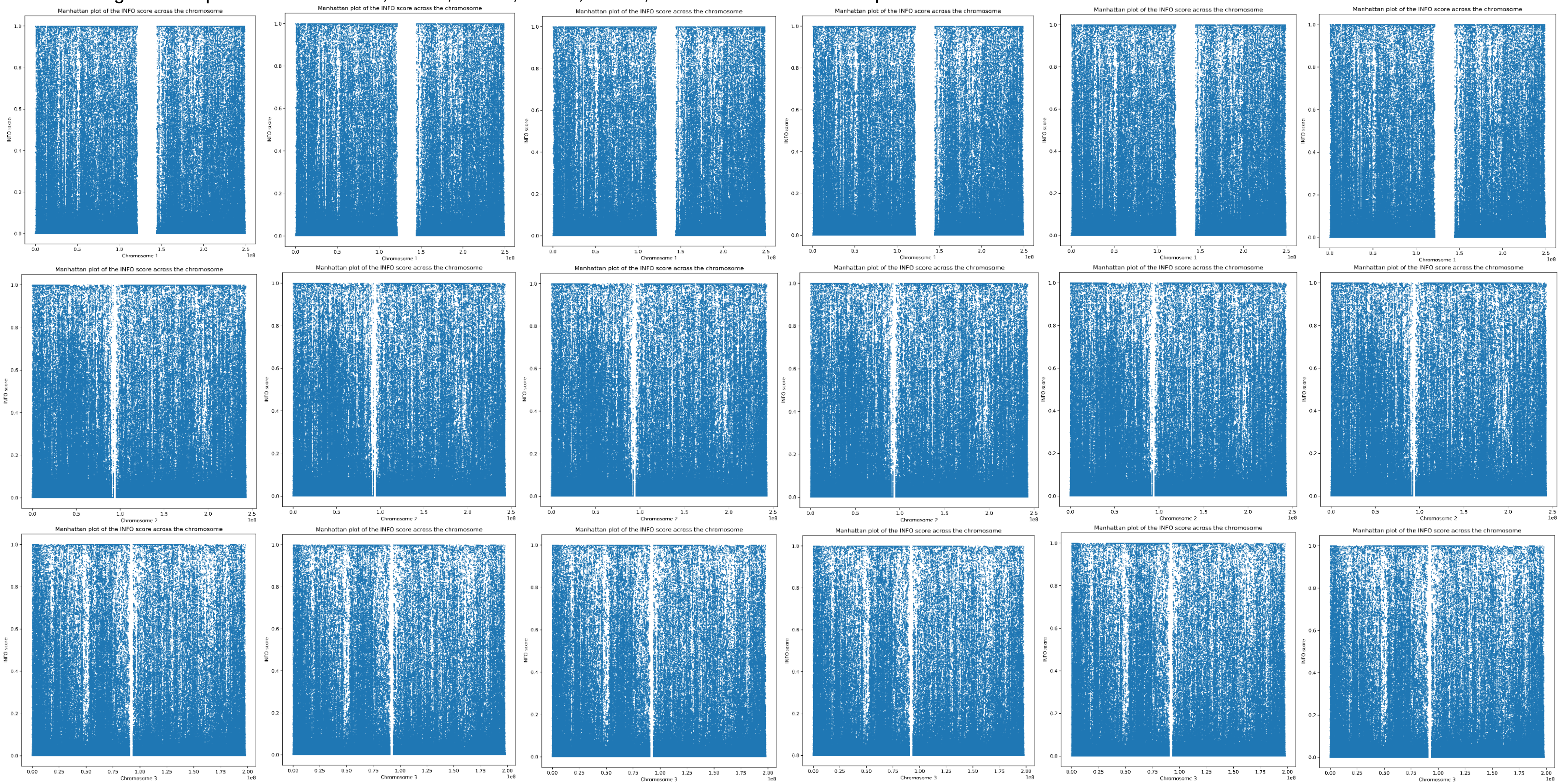

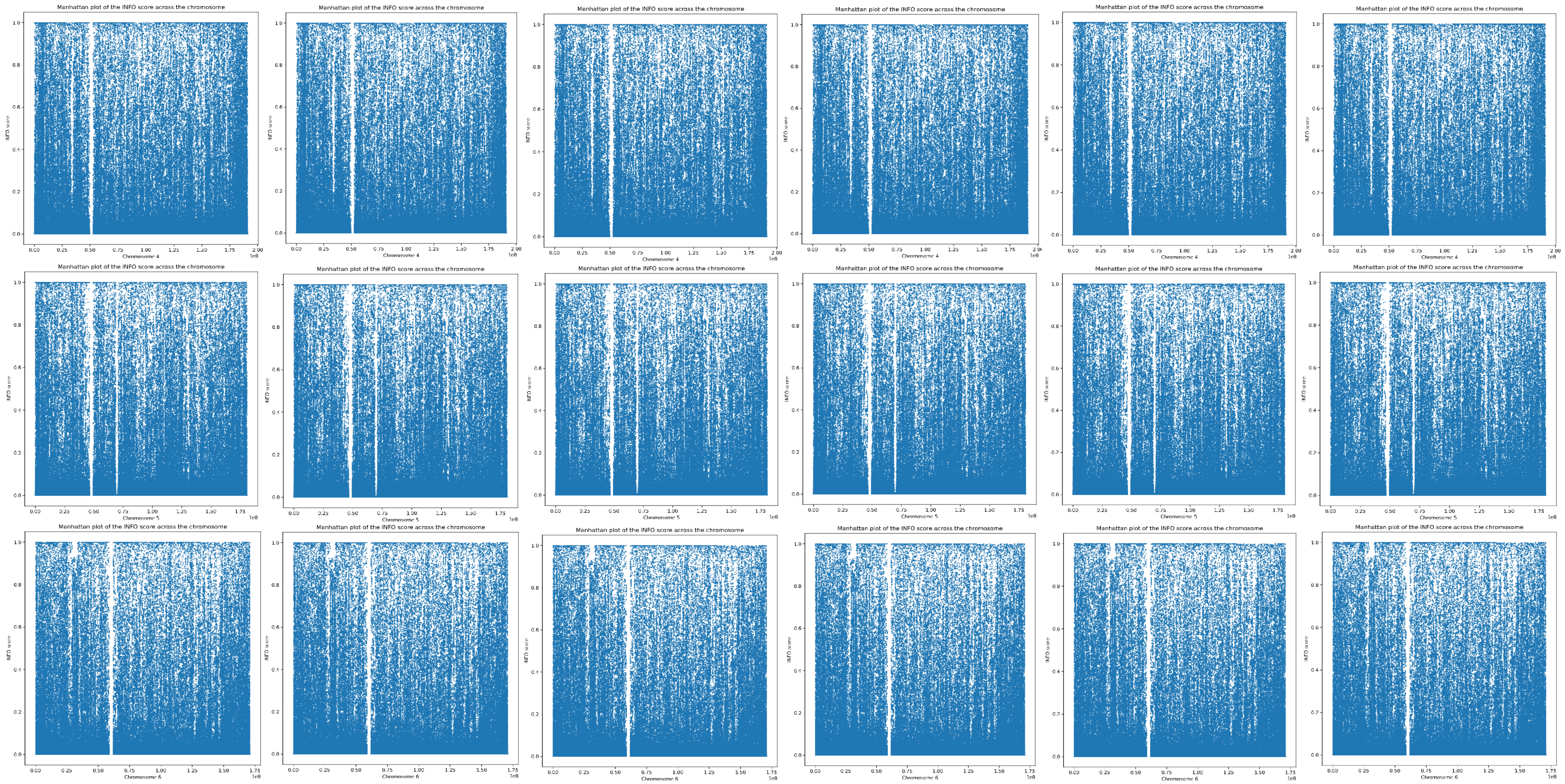
Manhattan plot of the INFO score across the chromosome
INFO score
Chromosome 4
Chromosome 5
Chromosome 6
1e8

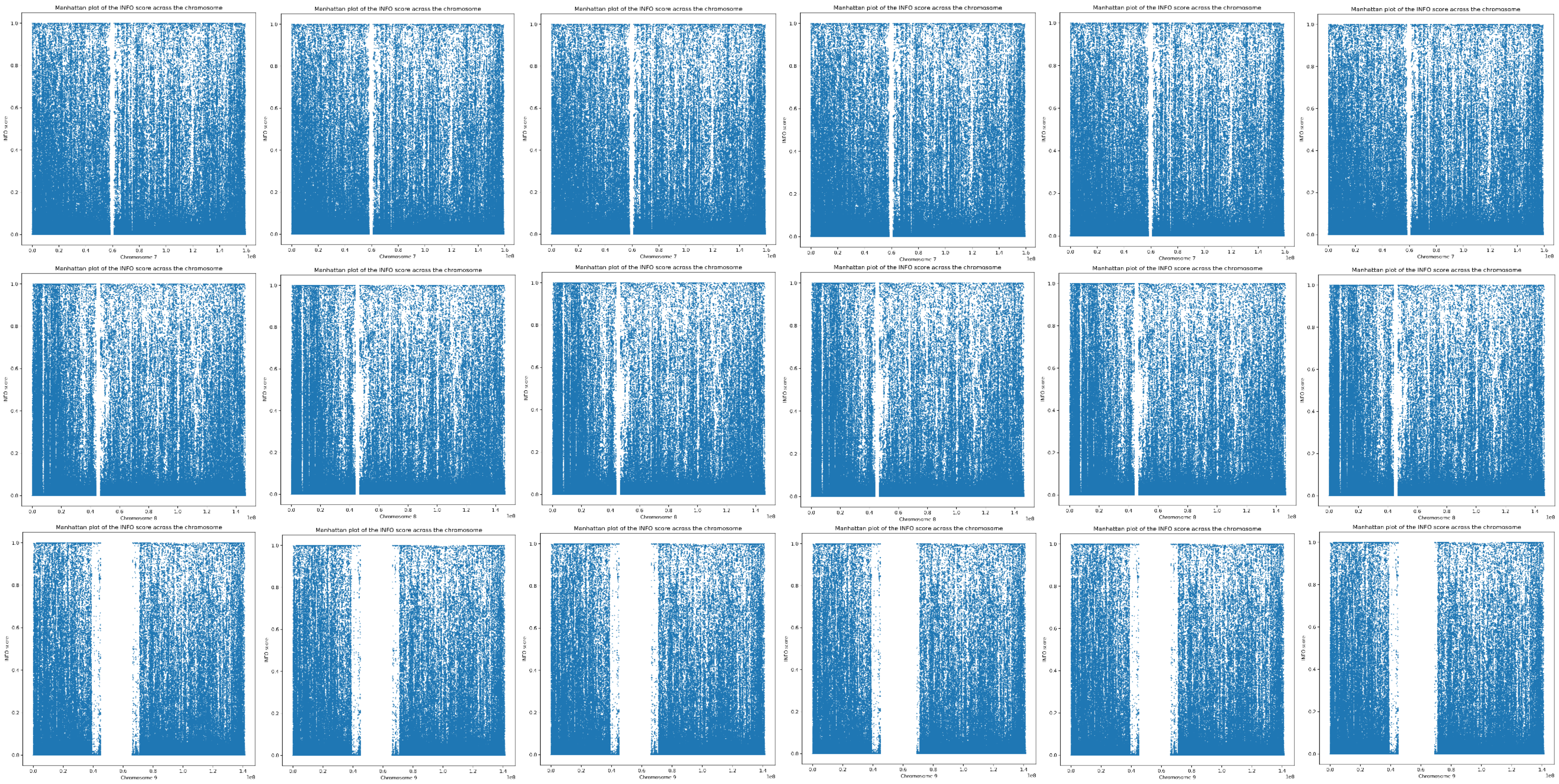
Manhattan plot of the INFO score across the chromosome
INFO score
Chromosome 7
Chromosome 8
Chromosome 9
1e8

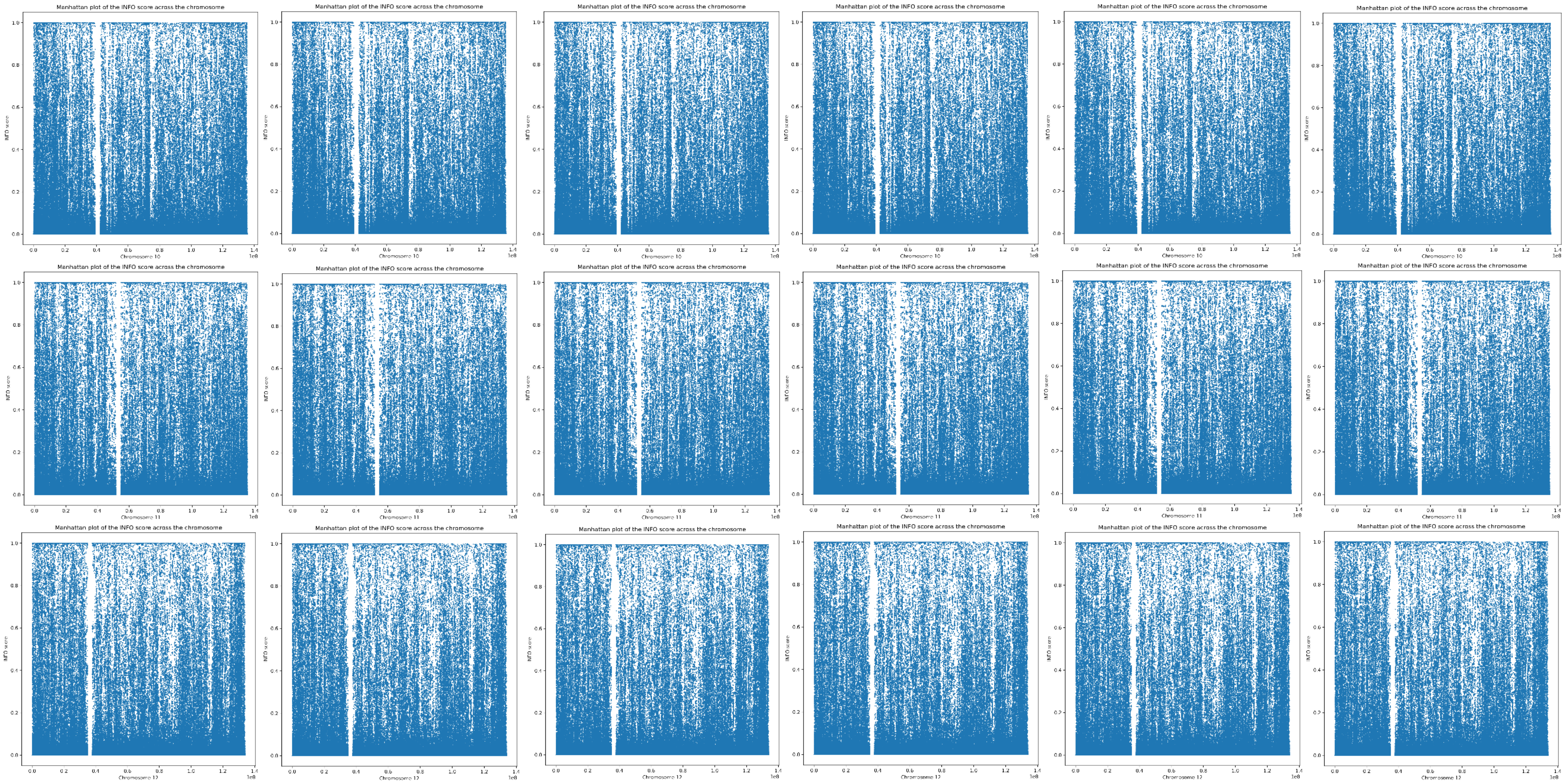
Manhattan plot of the INFO score across the chromosome
INFO score
Chromosome 10
Chromosome 11
Chromosome 12
1e8

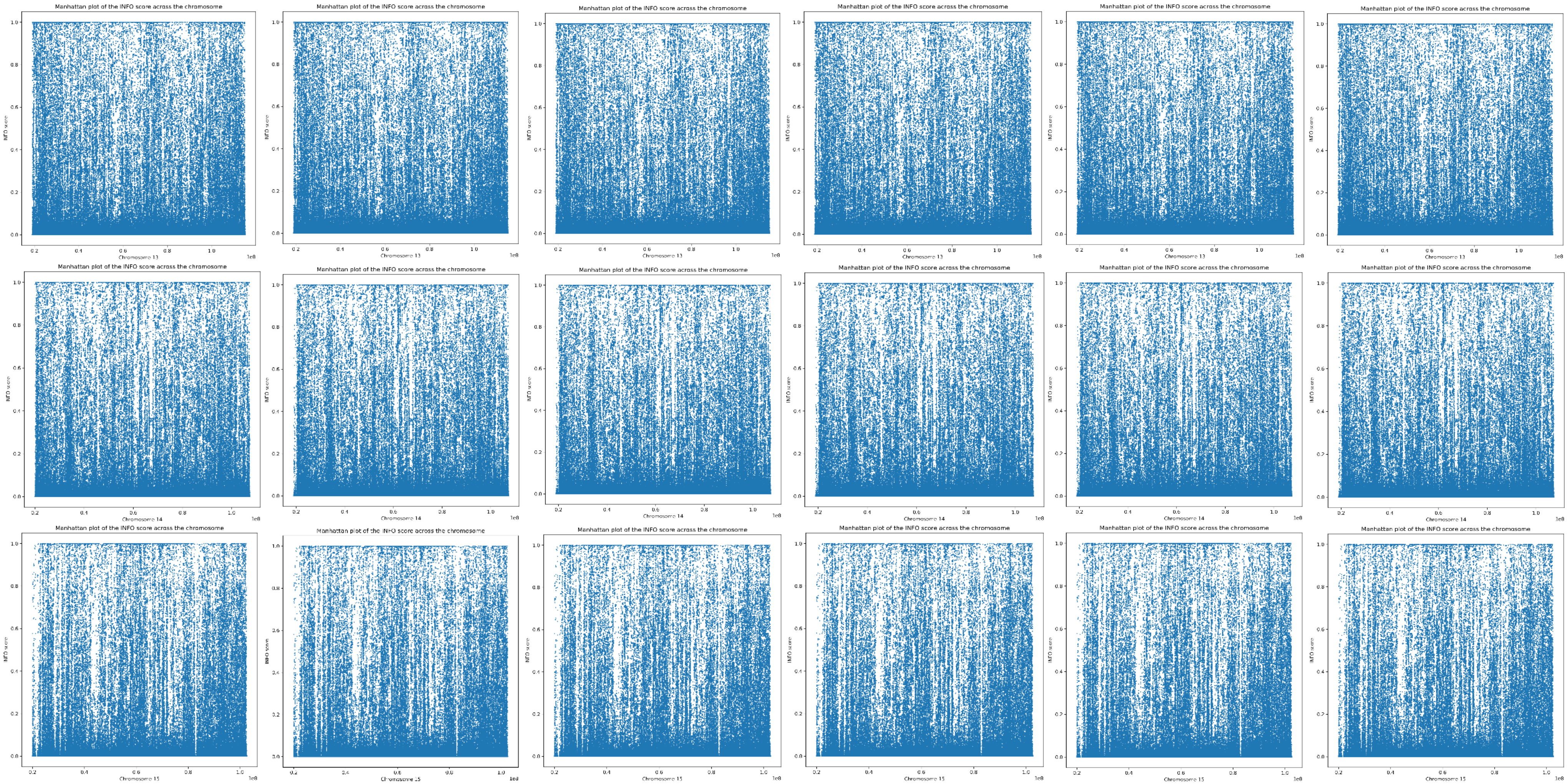

Manhattan plot of the INFO score across the chromosome
INFO score
1.0
0.8
0.6
0.4
0.2
0.0
0.2
0.4
0.6
0.8
1.0
1e8
Chromosome 13
Manhattan plot of the INFO score across the chromosome
INFO score
Chromosome 14
Manhattan plot of the INFO score across the chromosome
INFO score
Chromosome 15

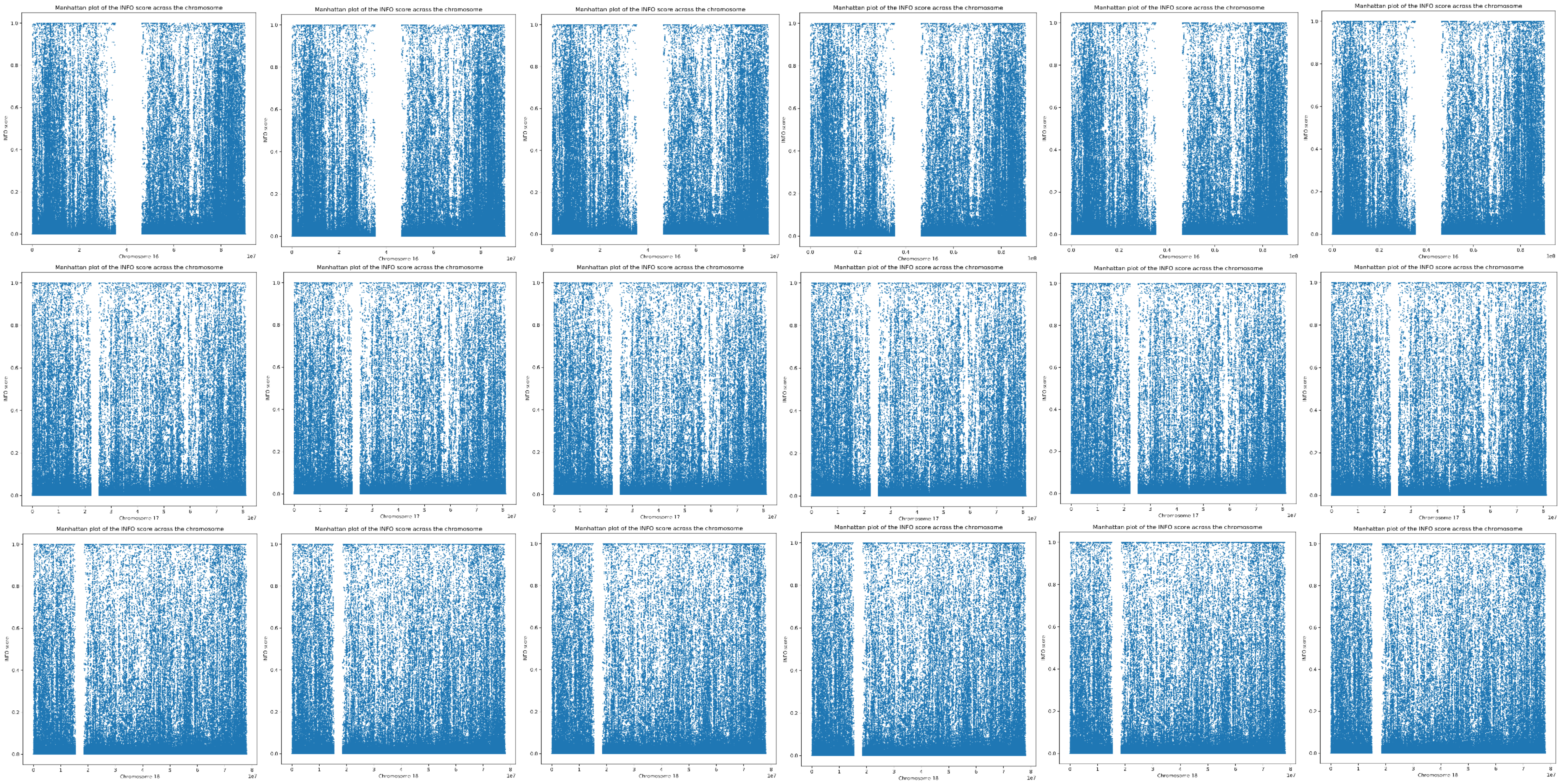
Manhattan plot of the INFO score across the chromosome
INFO score
Chromosome 16
Chromosome 17
Chromosome 18

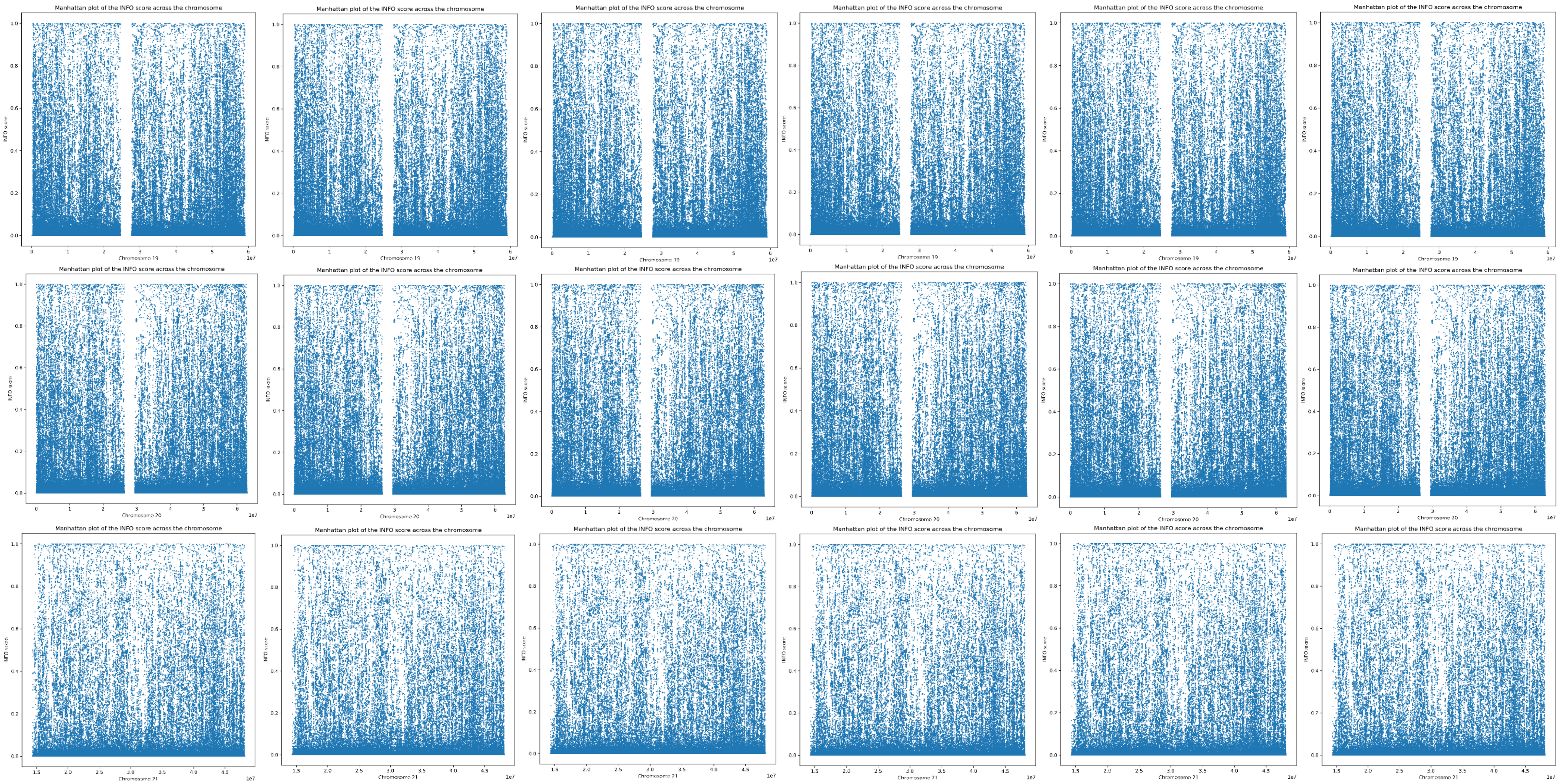

Manhattan plot of the INFO score across the chromosome
INFO score
Chromosome 19
Chromosome 20
Chromosome 21
1e7

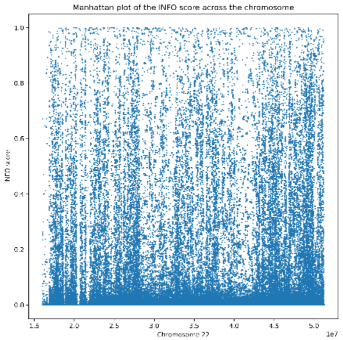
Manhattan plot of the INFO score across the chromosome
INFO score
Chromosome 22
1e7

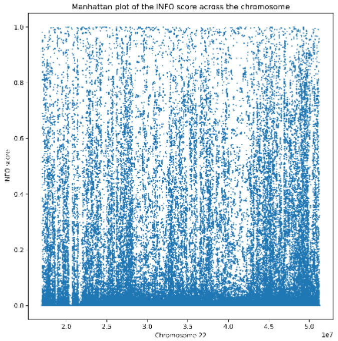
Manhattan plot of the INFO score across the chromosome
INFO score
Chromosome 22
1e7

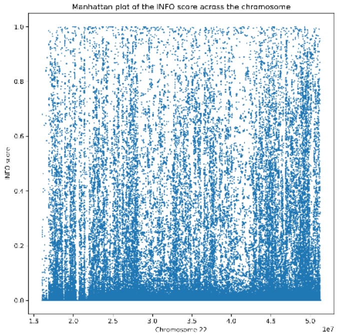
Manhattan plot of the INFO score across the chromosome
INFO score
Chromosome 22
1e7

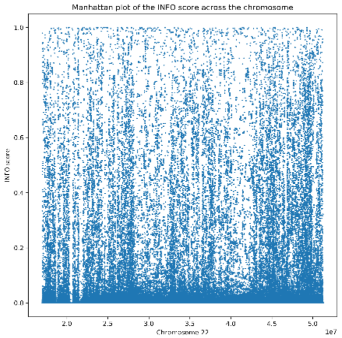
Manhattan plot of the INFO score across the chromosome
INFO score
Chromosome 22
1e7

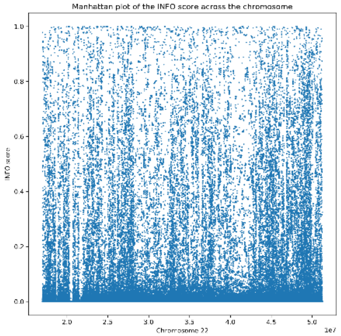
Manhattan plot of the INFO score across the chromosome
INFO score
Chromosome 22
1e7

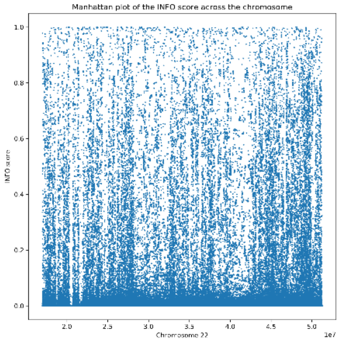
Manhattan plot of the INFO score across the chromosome
INFO score
Chromosome 22
1e7

Supplementary Figure 4. Imputation quality plots showing minor allele frequency (MAF) in MoBa imputed data vs imputation quality scores (INFO) for each chromosome and imputation batch. Plots from left to right are Imputation batch HCE, OMNI, GSA1, GSA2, GSA3, and GSA4. Plots from top to bottom are chromosome 1-22.

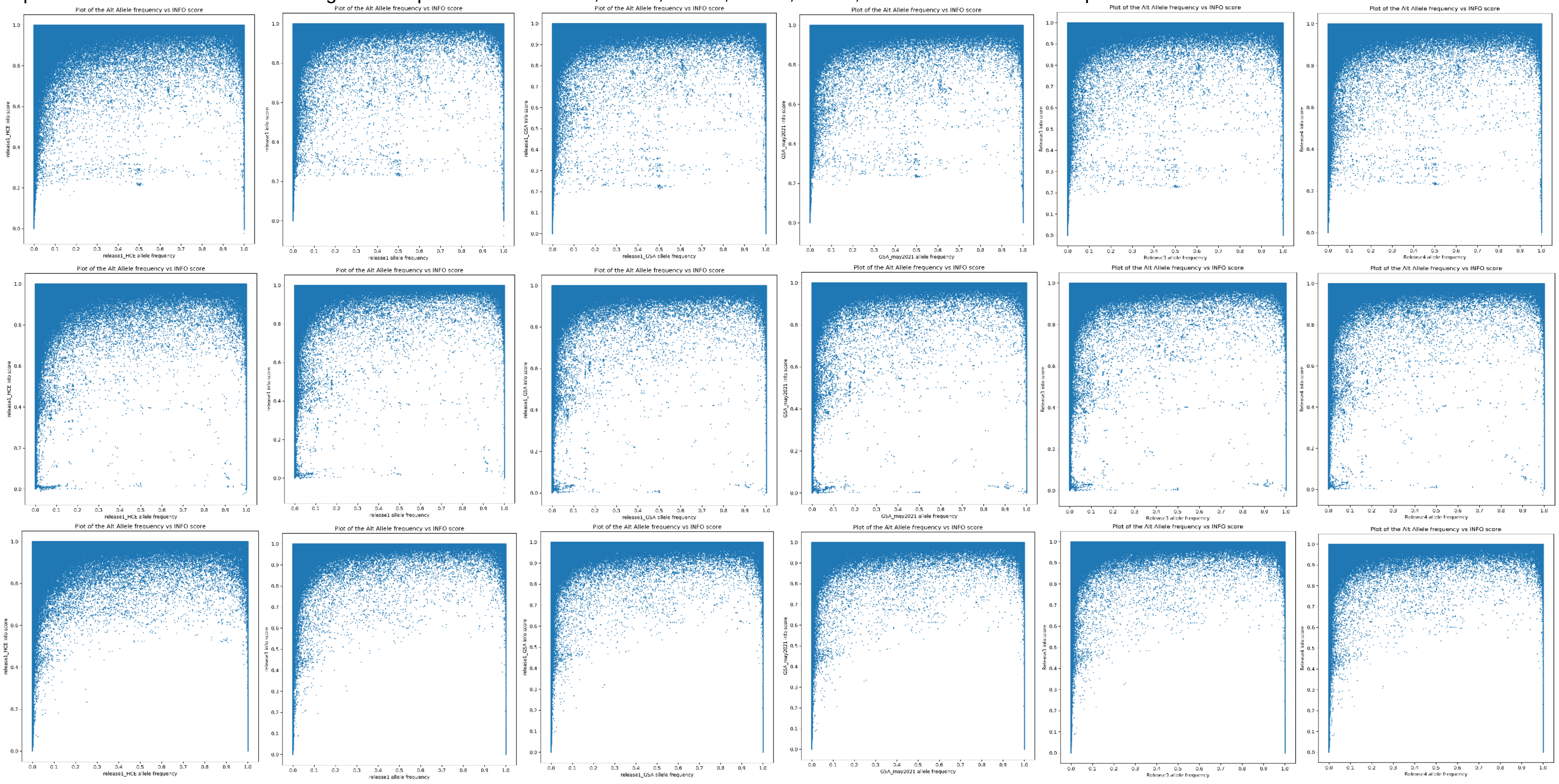

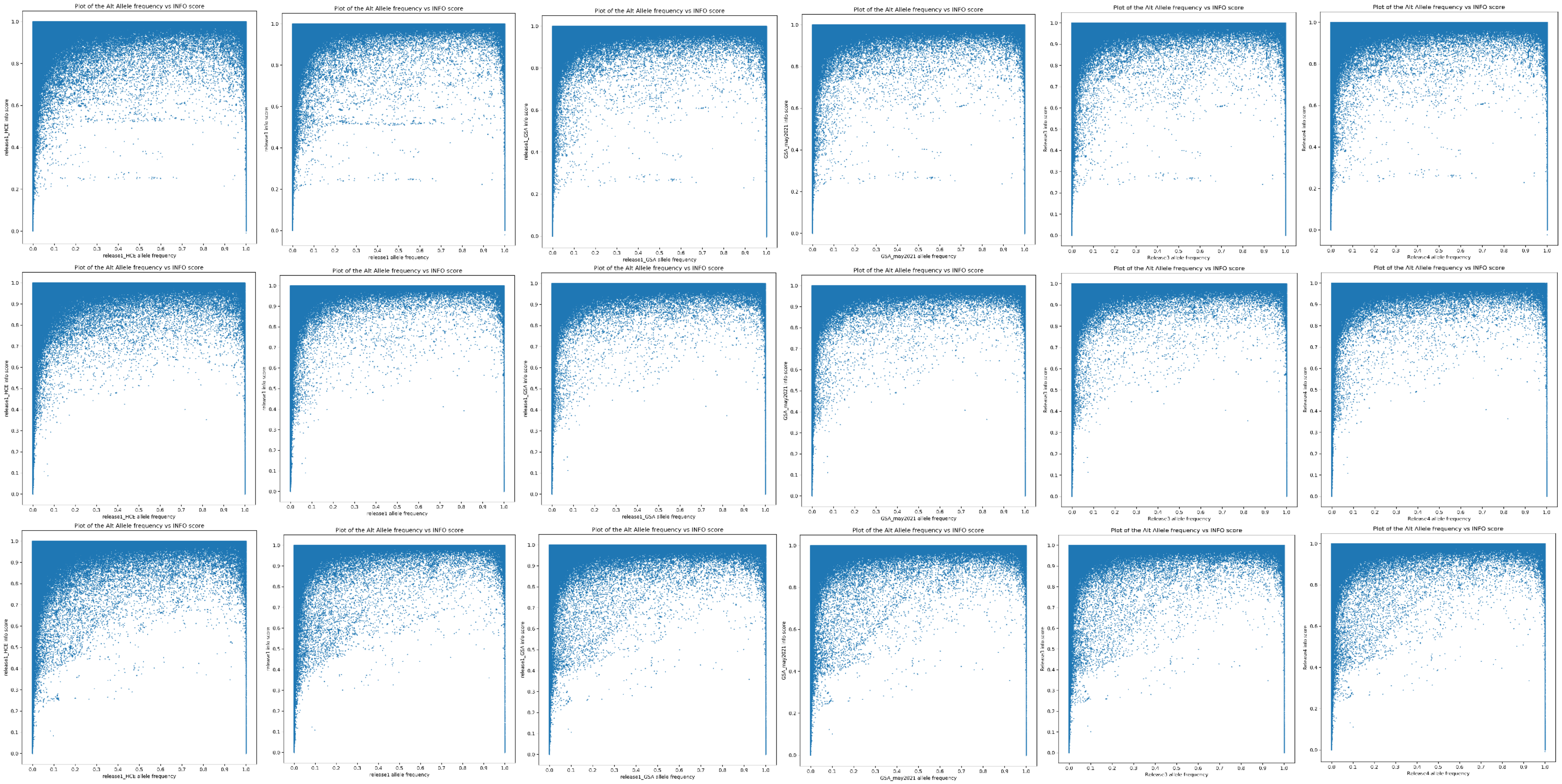
Plot of the Alt Allele frequency vs INFO score
release1_HCE info score
release1_HCE allele frequency
Plot of the Alt Allele frequency vs INFO score
release1 info score
release1 allele frequency
Plot of the Alt Allele frequency vs INFO score
release1_GSA info score
release1_GSA allele frequency
Plot of the Alt Allele frequency vs INFO score
GSA_may2021 info score
GSA_may2021 allele frequency
Plot of the Alt Allele frequency vs INFO score
Release3 info score
Release3 allele frequency
Plot of the Alt Allele frequency vs INFO score
Release4 info score
Release4 allele frequency
Plot of the Alt Allele frequency vs INFO score
release1_HCE info score
release1_HCE allele frequency
Plot of the Alt Allele frequency vs INFO score
release1 info score
release1 allele frequency
Plot of the Alt Allele frequency vs INFO score
release1_GSA info score
release1_GSA allele frequency
Plot of the Alt Allele frequency vs INFO score
GSA_may2021 info score
GSA_may2021 allele frequency
Plot of the Alt Allele frequency vs INFO score
Release3 info score
Release3 allele frequency
Plot of the Alt Allele frequency vs INFO score
Release4 info score
Release4 allele frequency
Plot of the Alt Allele frequency vs INFO score
release1_HCE info score
release1_HCE allele frequency
Plot of the Alt Allele frequency vs INFO score
release1 info score
release1 allele frequency
Plot of the Alt Allele frequency vs INFO score
release1_GSA info score
release1_GSA allele frequency
Plot of the Alt Allele frequency vs INFO score
GSA_may2021 info score
GSA_may2021 allele frequency
Plot of the Alt Allele frequency vs INFO score
Release3 info score
Release3 allele frequency
Plot of the Alt Allele frequency vs INFO score
Release4 info score
Release4 allele frequency

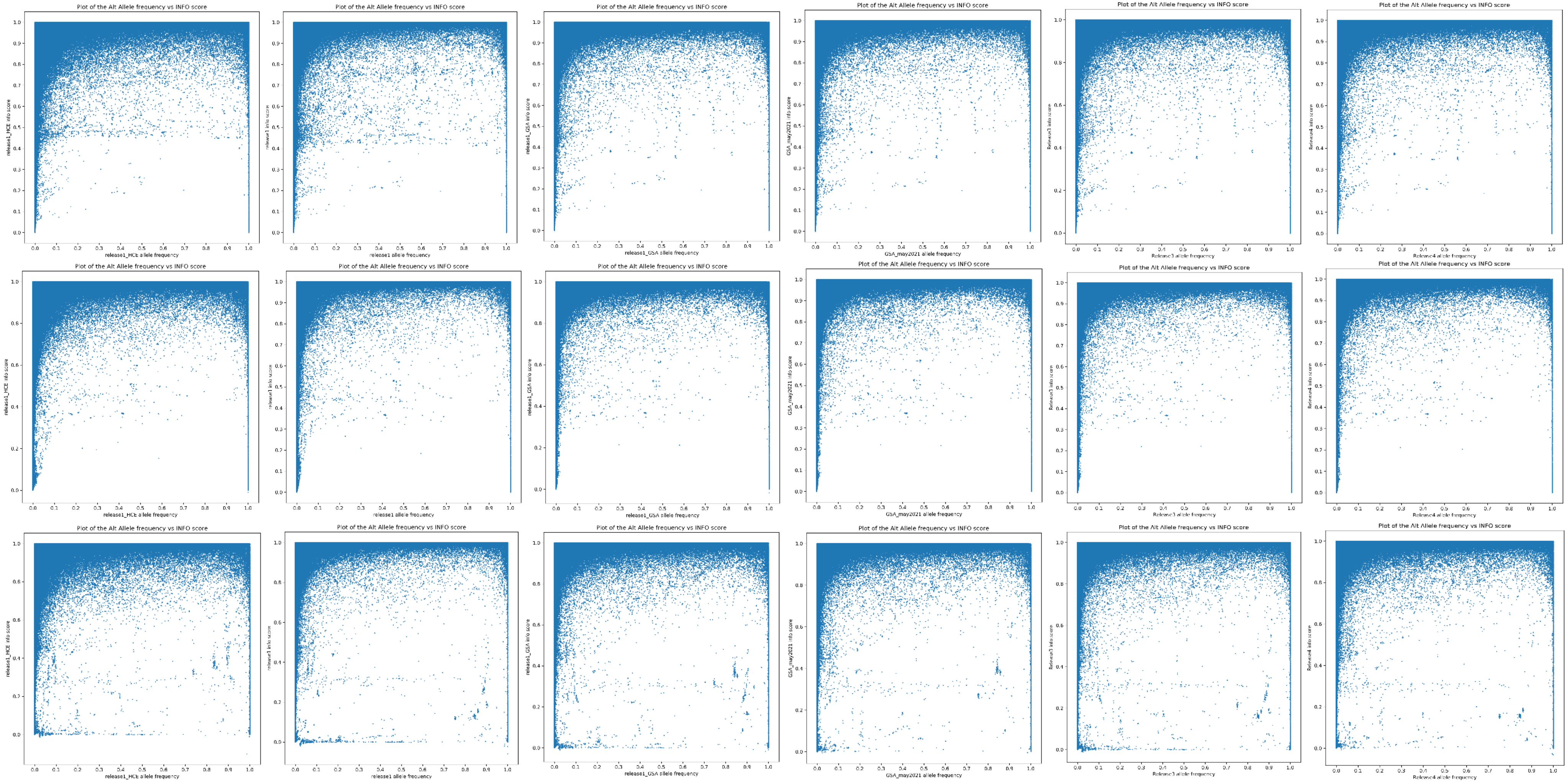
Plot of the Alt Allele frequency vs INFO score
release1_HCE info score
release1_HCE allele frequency
release1 info score
release1 allele frequency
release1_GSA info score
release1_GSA allele frequency
GSA_may2021 info score
GSA_may2021 allele frequency
Release3 info score
Release3 allele frequency
Release4 info score
Release4 allele frequency

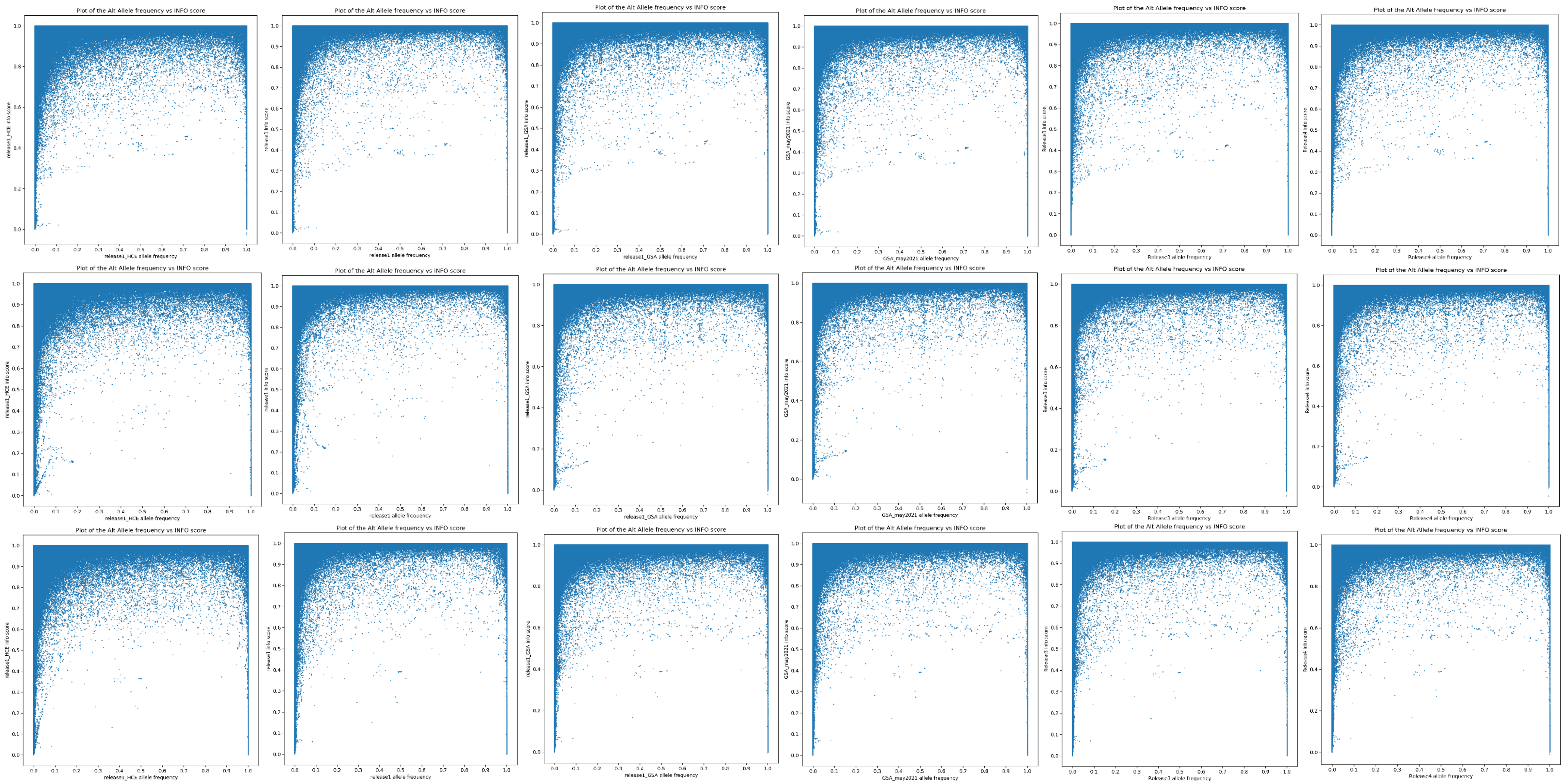
Plot of the Alt Allele frequency vs INFO score
release1_HCE info score
release1_HCE allele frequency
release1 info score
release1 allele frequency
release1_GSA info score
release1_GSA allele frequency
GSA_may2021 info score
GSA_may2021 allele frequency
Release3 info score
Release3 allele frequency
Release4 info score
Release4 allele frequency

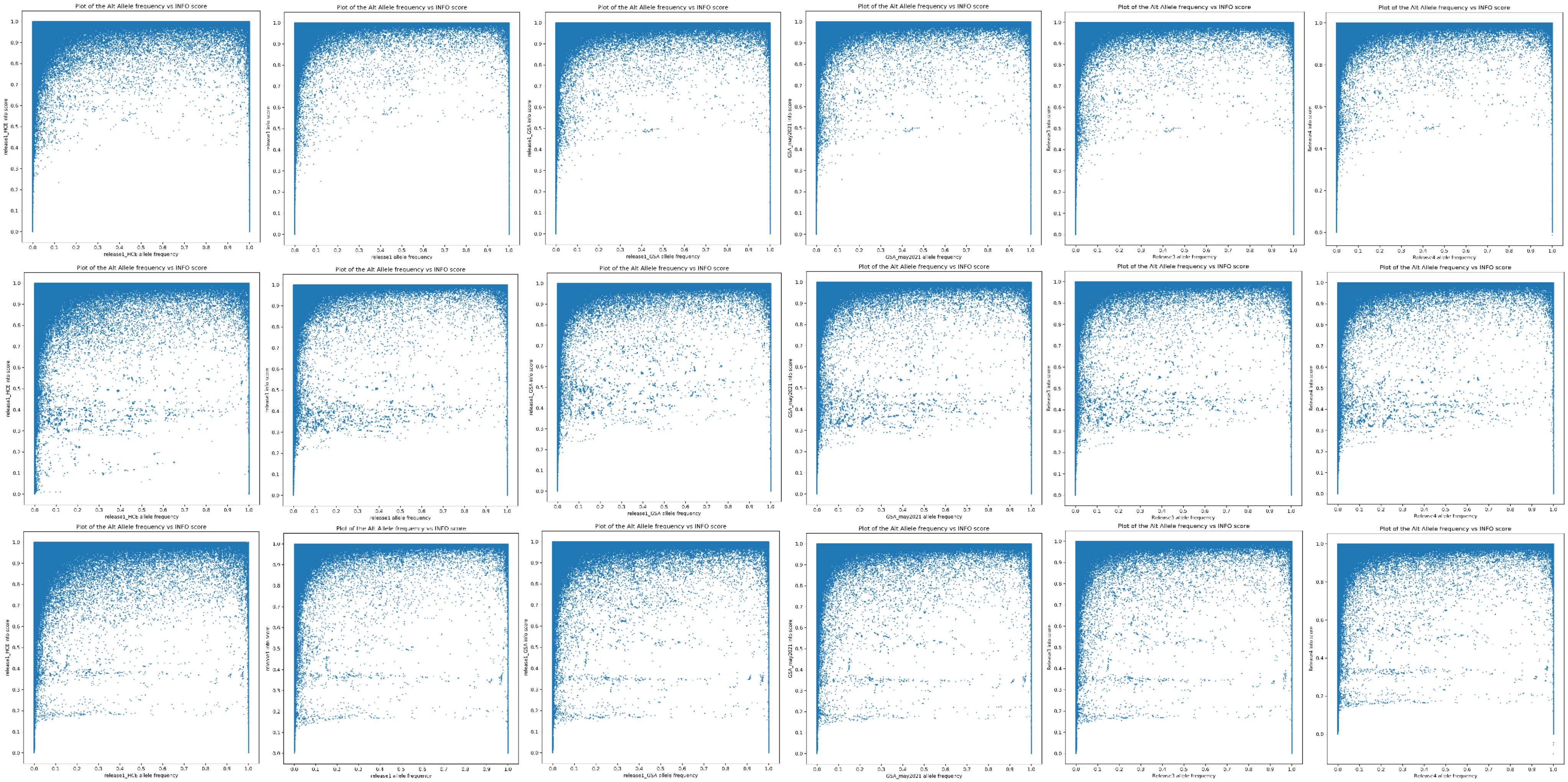
Plot of the Alt Allele frequency vs INFO score
release1_HCE allele frequency
release1_HCE info score
release1 allele frequency
release1 info score
release1_GSA allele frequency
release1_GSA info score
GSA_may2021 allele frequency
GSA_may2021 info score
Release3 allele frequency
Release3 info score
Release4 allele frequency
Release4 info score

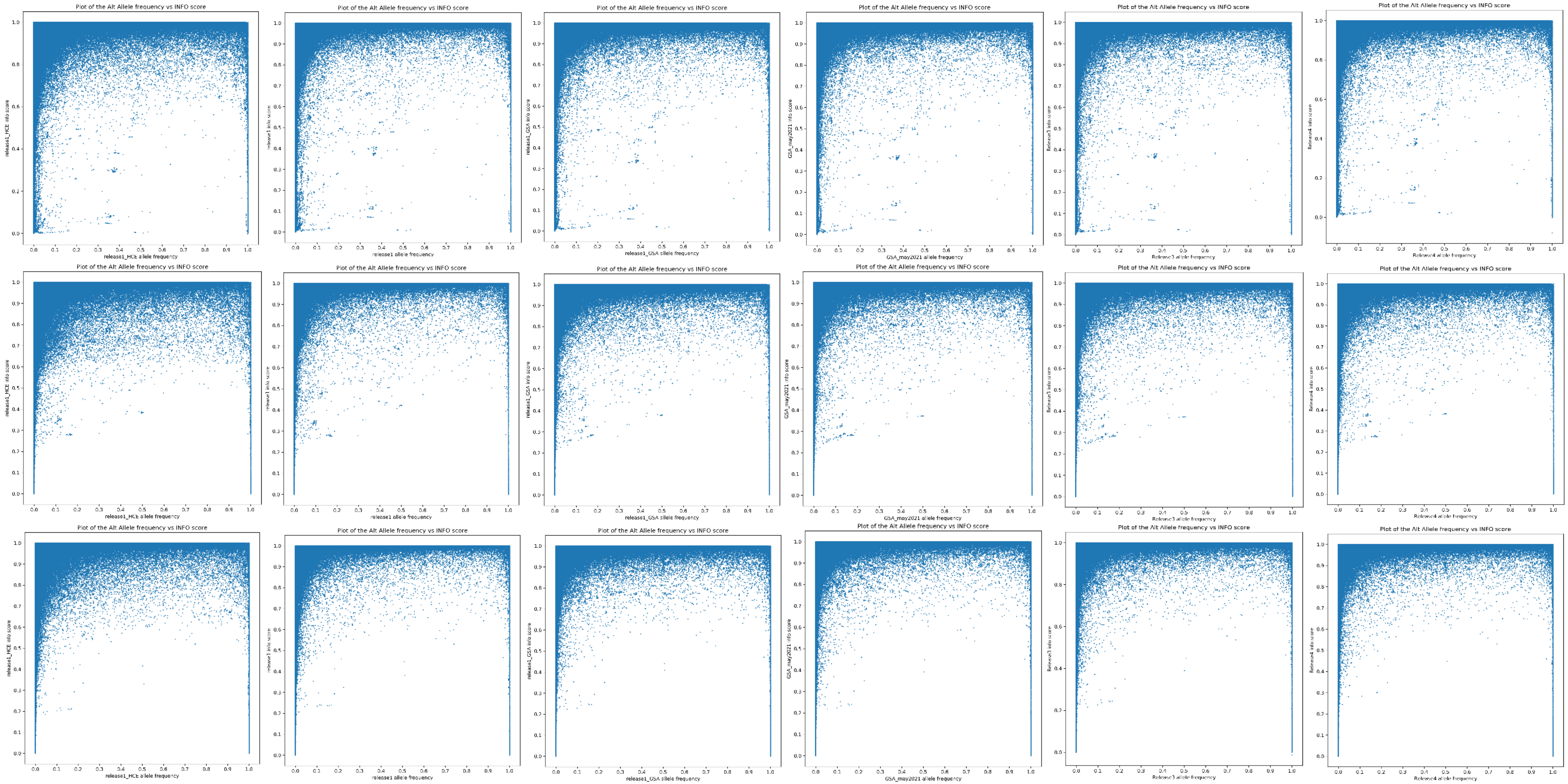
Plot of the Alt Allele frequency vs INFO score
release1_HCE info score
release1_HCE allele frequency
Plot of the Alt Allele frequency vs INFO score
release1 info score
release1 allele frequency
Plot of the Alt Allele frequency vs INFO score
release1_GSA info score
release1_GSA allele frequency
Plot of the Alt Allele frequency vs INFO score
GSA_may2021 info score
GSA_may2021 allele frequency
Plot of the Alt Allele frequency vs INFO score
Release3 info score
Release3 allele frequency
Plot of the Alt Allele frequency vs INFO score
Release4 info score
Release4 allele frequency
Plot of the Alt Allele frequency vs INFO score
release1_HCE info score
release1_HCE allele frequency
Plot of the Alt Allele frequency vs INFO score
release1 info score
release1 allele frequency
Plot of the Alt Allele frequency vs INFO score
release1_GSA info score
release1_GSA allele frequency
Plot of the Alt Allele frequency vs INFO score
GSA_may2021 info score
GSA_may2021 allele frequency
Plot of the Alt Allele frequency vs INFO score
Release3 info score
Release3 allele frequency
Plot of the Alt Allele frequency vs INFO score
Release4 info score
Release4 allele frequency
Plot of the Alt Allele frequency vs INFO score
release1_HCE info score
release1_HCE allele frequency
Plot of the Alt Allele frequency vs INFO score
release1 info score
release1 allele frequency
Plot of the Alt Allele frequency vs INFO score
release1_GSA info score
release1_GSA allele frequency
Plot of the Alt Allele frequency vs INFO score
GSA_may2021 info score
GSA_may2021 allele frequency
Plot of the Alt Allele frequency vs INFO score
Release3 info score
Release3 allele frequency
Plot of the Alt Allele frequency vs INFO score
Release4 info score
Release4 allele frequency

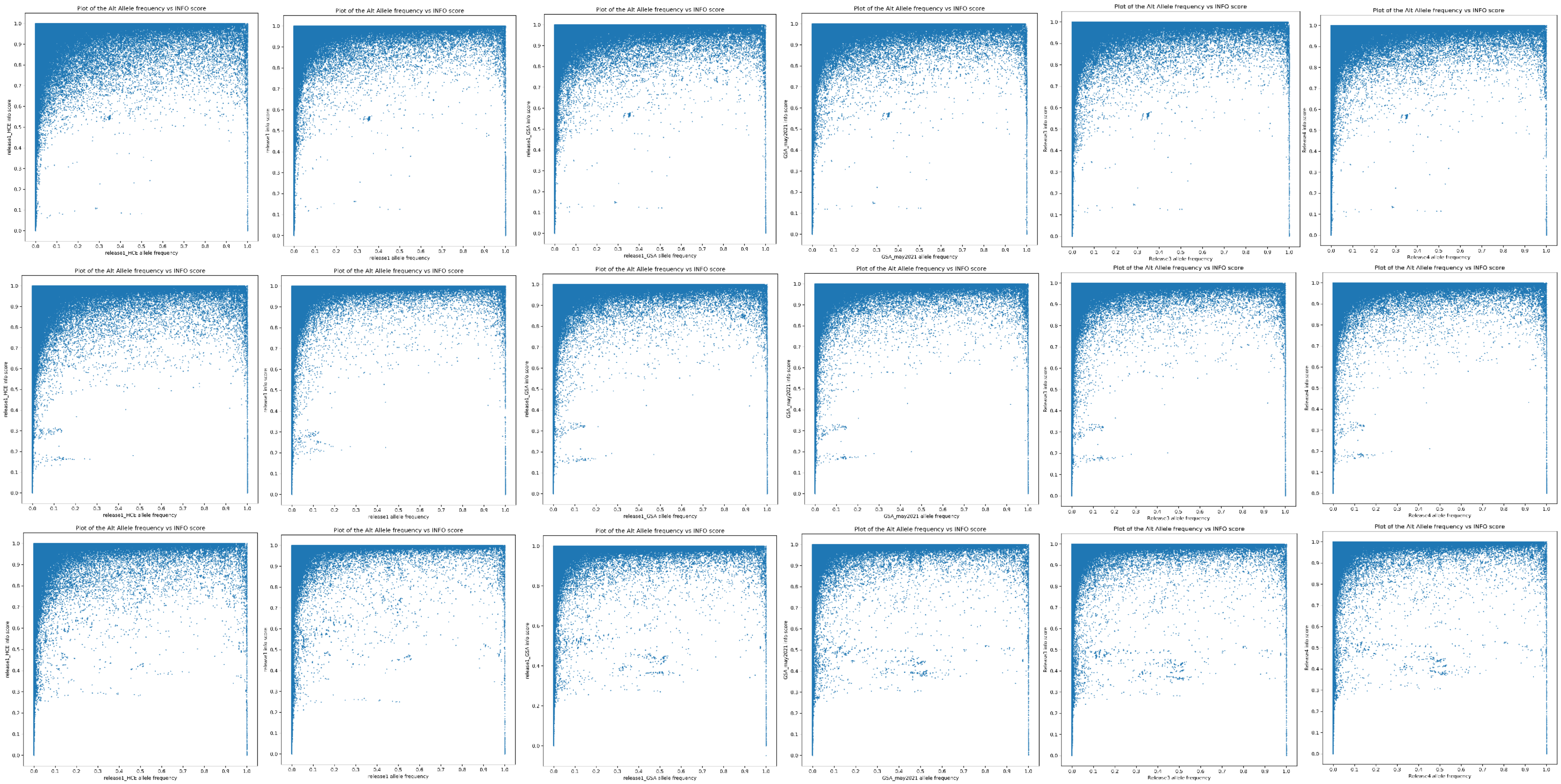
Plot of the Alt Allele frequency vs INFO score
release1_HCE allele frequency
Plot of the Alt Allele frequency vs INFO score
release1 allele frequency
Plot of the Alt Allele frequency vs INFO score
release1_GSA allele frequency
Plot of the Alt Allele frequency vs INFO score
GSA_may2021 allele frequency
Plot of the Alt Allele frequency vs INFO score
Release3 allele frequency
Plot of the Alt Allele frequency vs INFO score
Release4 allele frequency
Plot of the Alt Allele frequency vs INFO score
release1_HCE allele frequency
Plot of the Alt Allele frequency vs INFO score
release1 allele frequency
Plot of the Alt Allele frequency vs INFO score
release1_GSA allele frequency
Plot of the Alt Allele frequency vs INFO score
GSA_may2021 allele frequency
Plot of the Alt Allele frequency vs INFO score
Release3 allele frequency
Plot of the Alt Allele frequency vs INFO score
Release4 allele frequency
Plot of the Alt Allele frequency vs INFO score
release1_HCE allele frequency
Plot of the Alt Allele frequency vs INFO score
release1 allele frequency
Plot of the Alt Allele frequency vs INFO score
release1_GSA allele frequency
Plot of the Alt Allele frequency vs INFO score
GSA_may2021 allele frequency
Plot of the Alt Allele frequency vs INFO score
Release3 allele frequency
Plot of the Alt Allele frequency vs INFO score
Release4 allele frequency

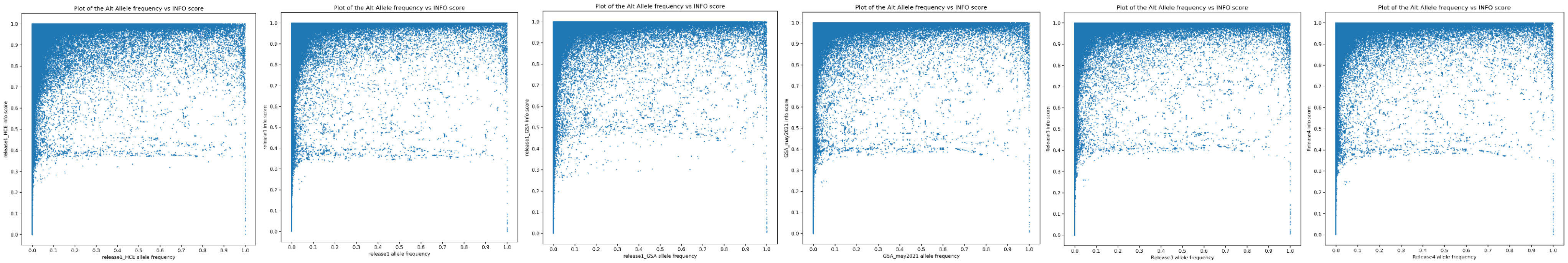
Plot of the Alt Allele frequency vs INFO score
release1_HCE info score
release1_HCE allele frequency
Plot of the Alt Allele frequency vs INFO score
release1 info score
release1 allele frequency
Plot of the Alt Allele frequency vs INFO score
release1_GSA info score
release1_GSA allele frequency
Plot of the Alt Allele frequency vs INFO score
GSA_may2021 info score
GSA_may2021 allele frequency
Plot of the Alt Allele frequency vs INFO score
Release3 info score
Release3 allele frequency
Plot of the Alt Allele frequency vs INFO score
Release4 info score
Release4 allele frequency

Supplementary Figure 5. Imputation quality plots showing minor allele frequency (MAF) in the Haplotype Reference Consortium vs MAF in MoBa imputed data for each chromosome and imputation batch. Plots from left to right are Imputation batch HCE, OMNI, GSA1, GSA2, GSA3, and GSA4. Plots from top to bottom are chromosome 1-22.

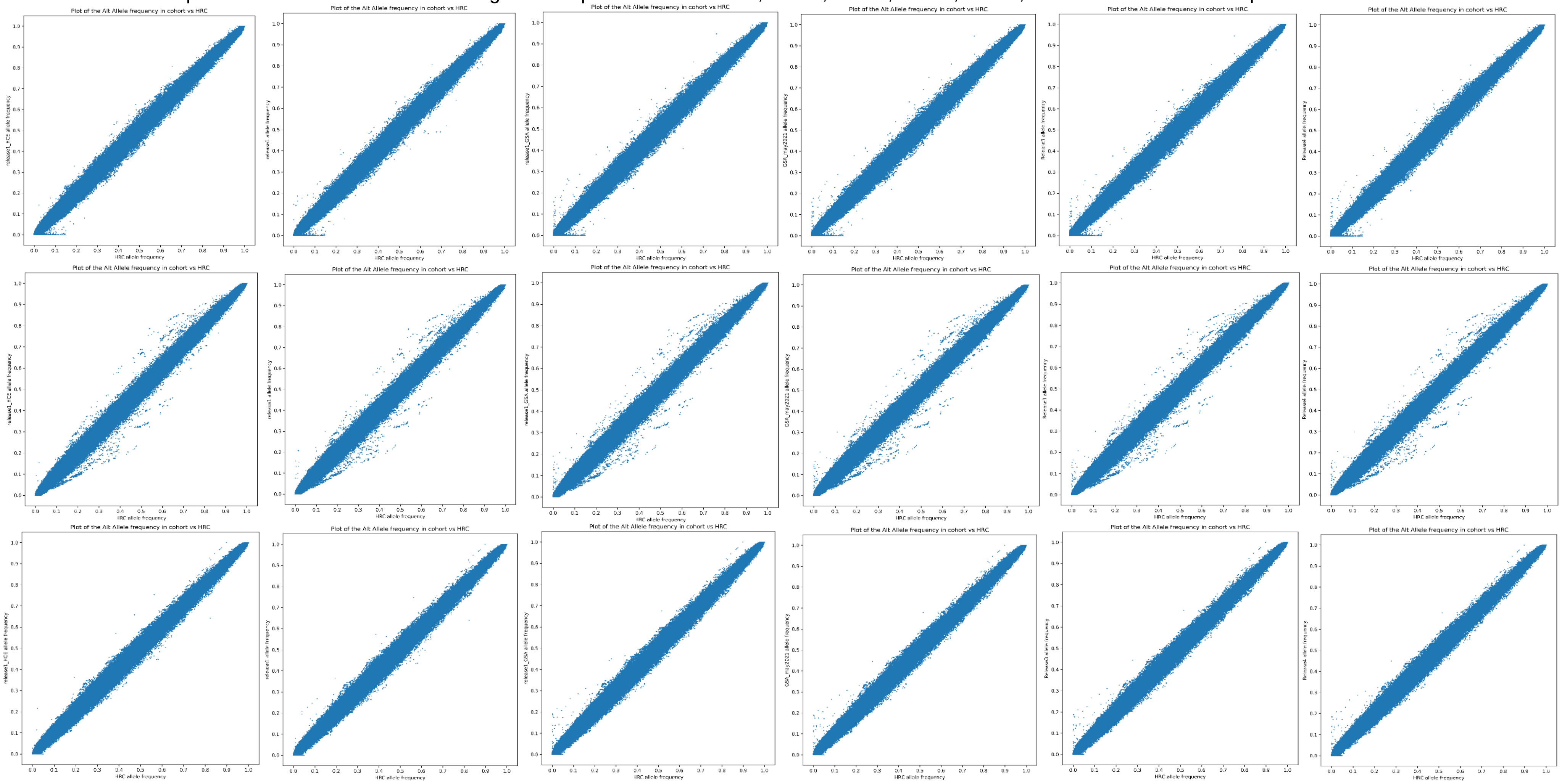

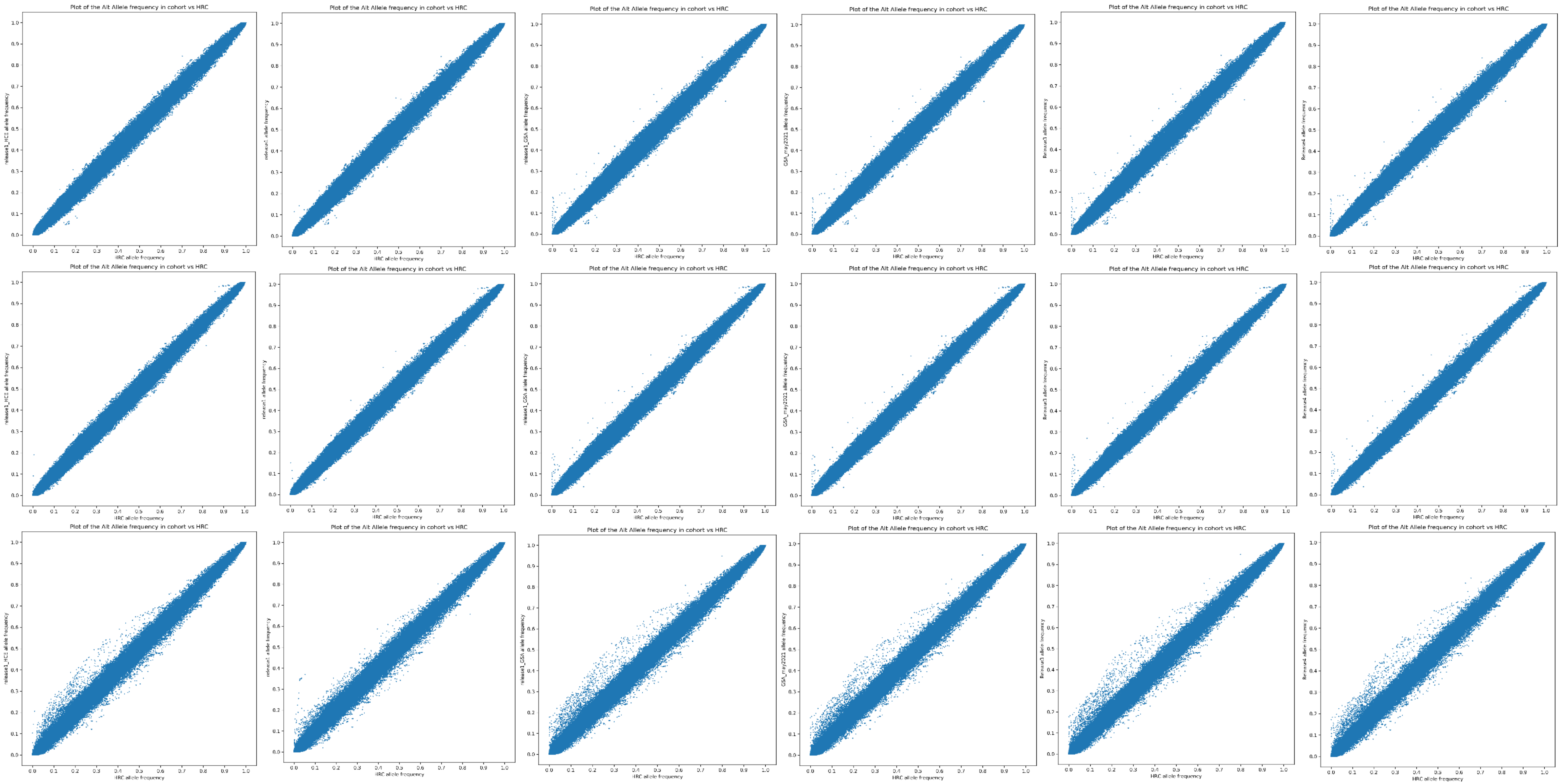
Plot of the Alt Allele frequency in cohort vs HRC
HRC allele frequency

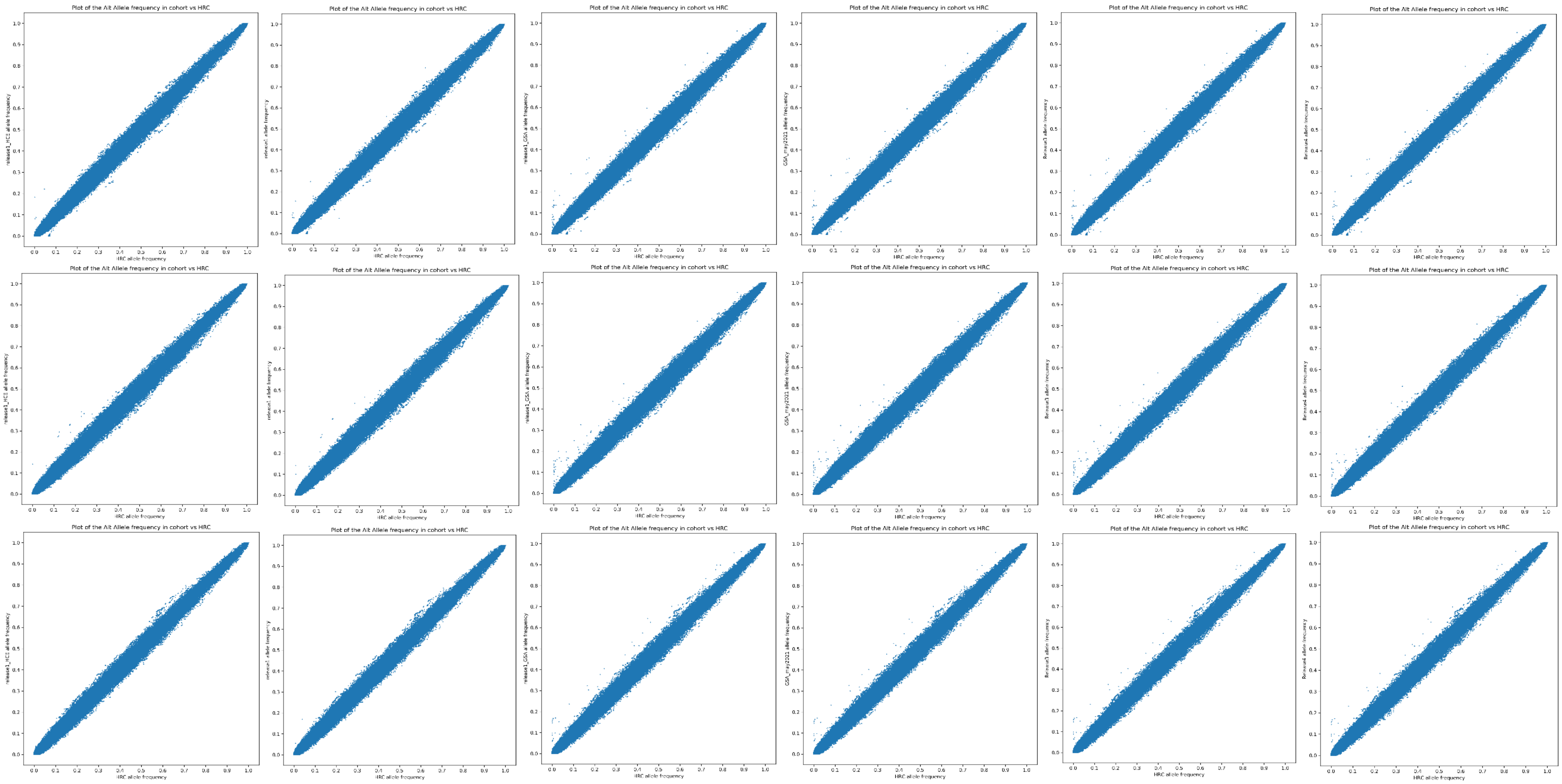
Plot of the Alt Allele frequency in cohort vs HRC
HRC allele frequency

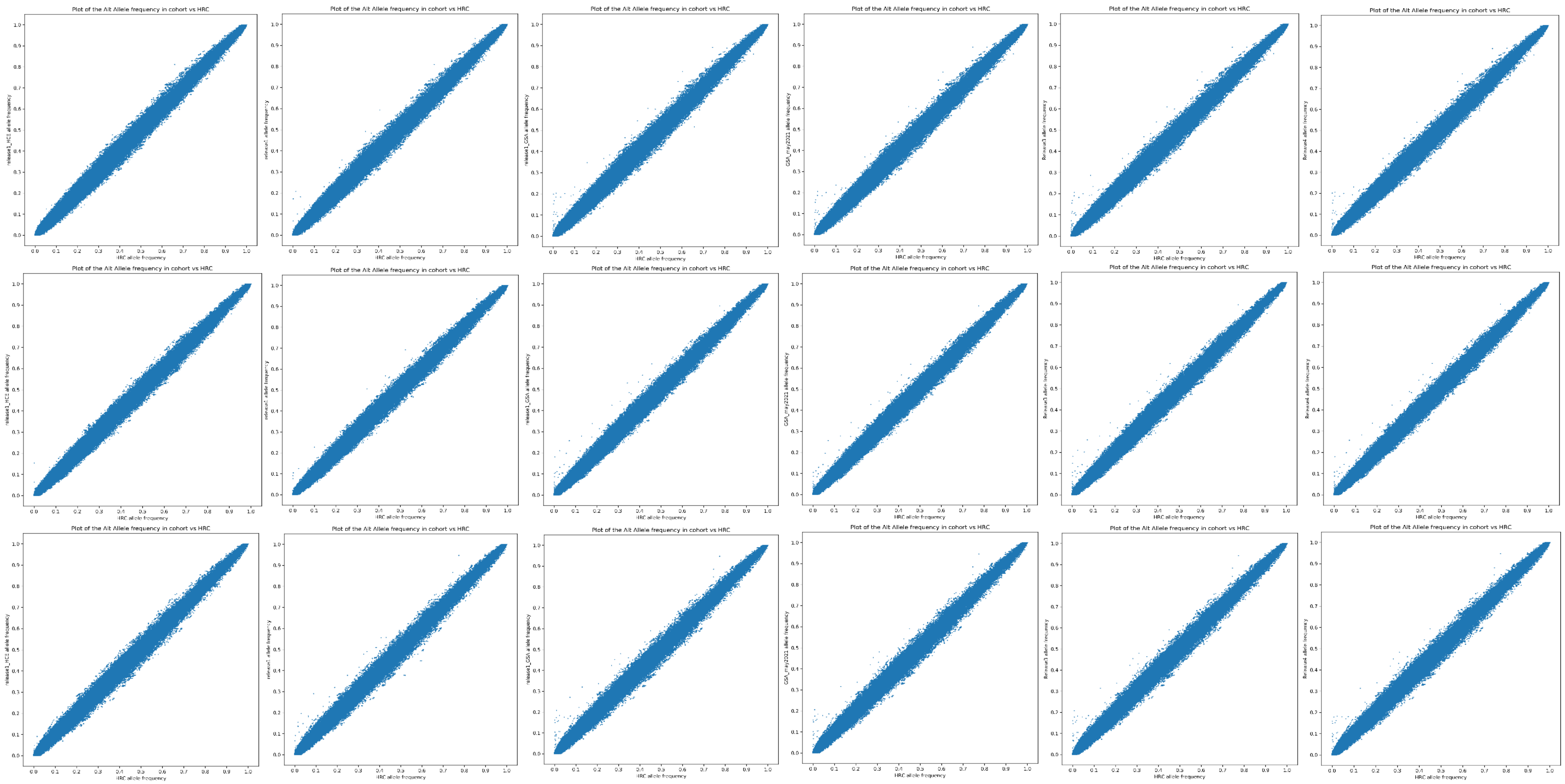
Plot of the Alt Allele frequency in cohort vs HRC
HRC allele frequency

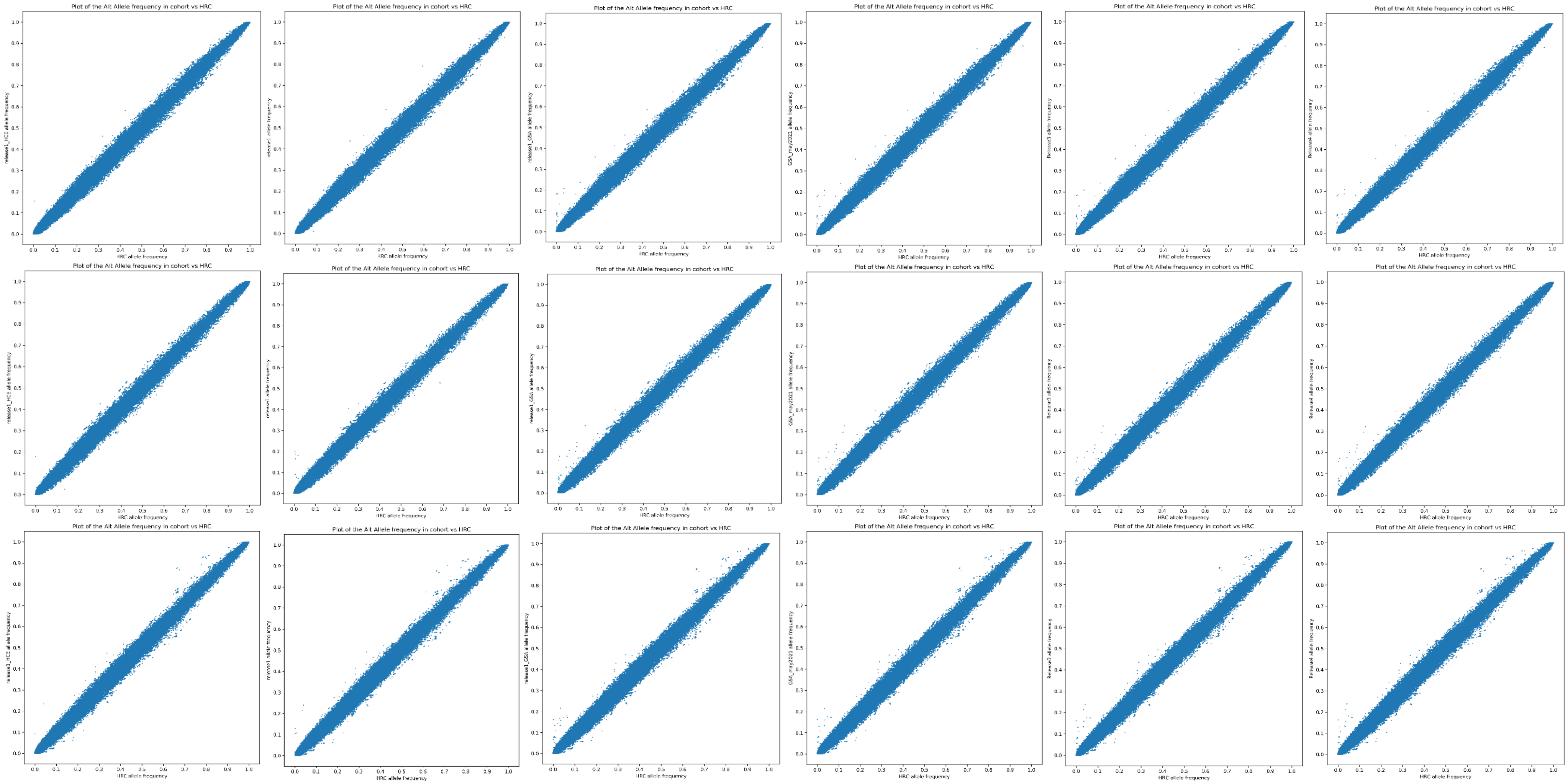
Plot of the Alt Allele frequency in cohort vs HRC
HRC allele frequency

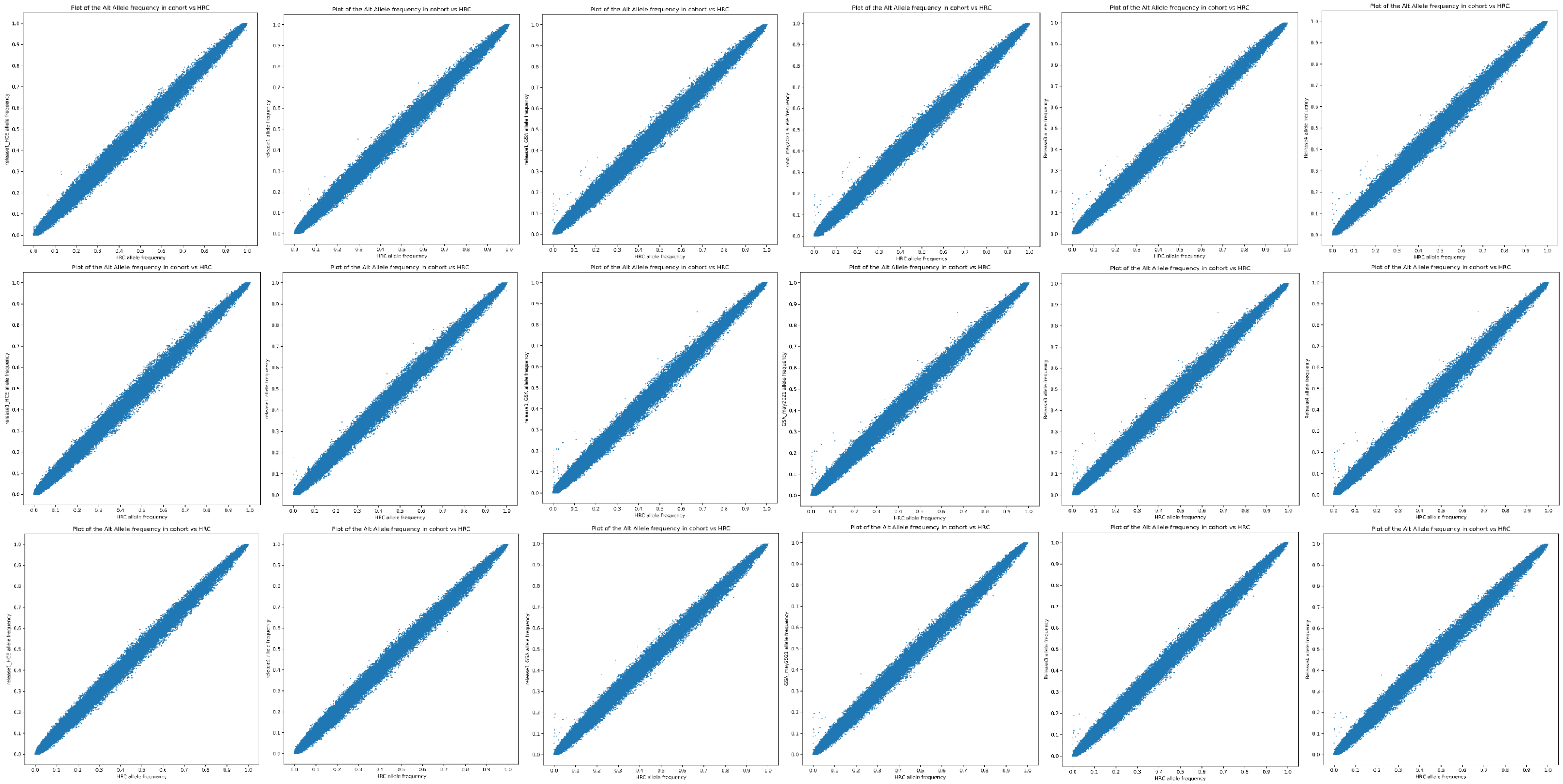
Plot of the Alt Allele frequency in cohort vs HRC
HRC allele frequency

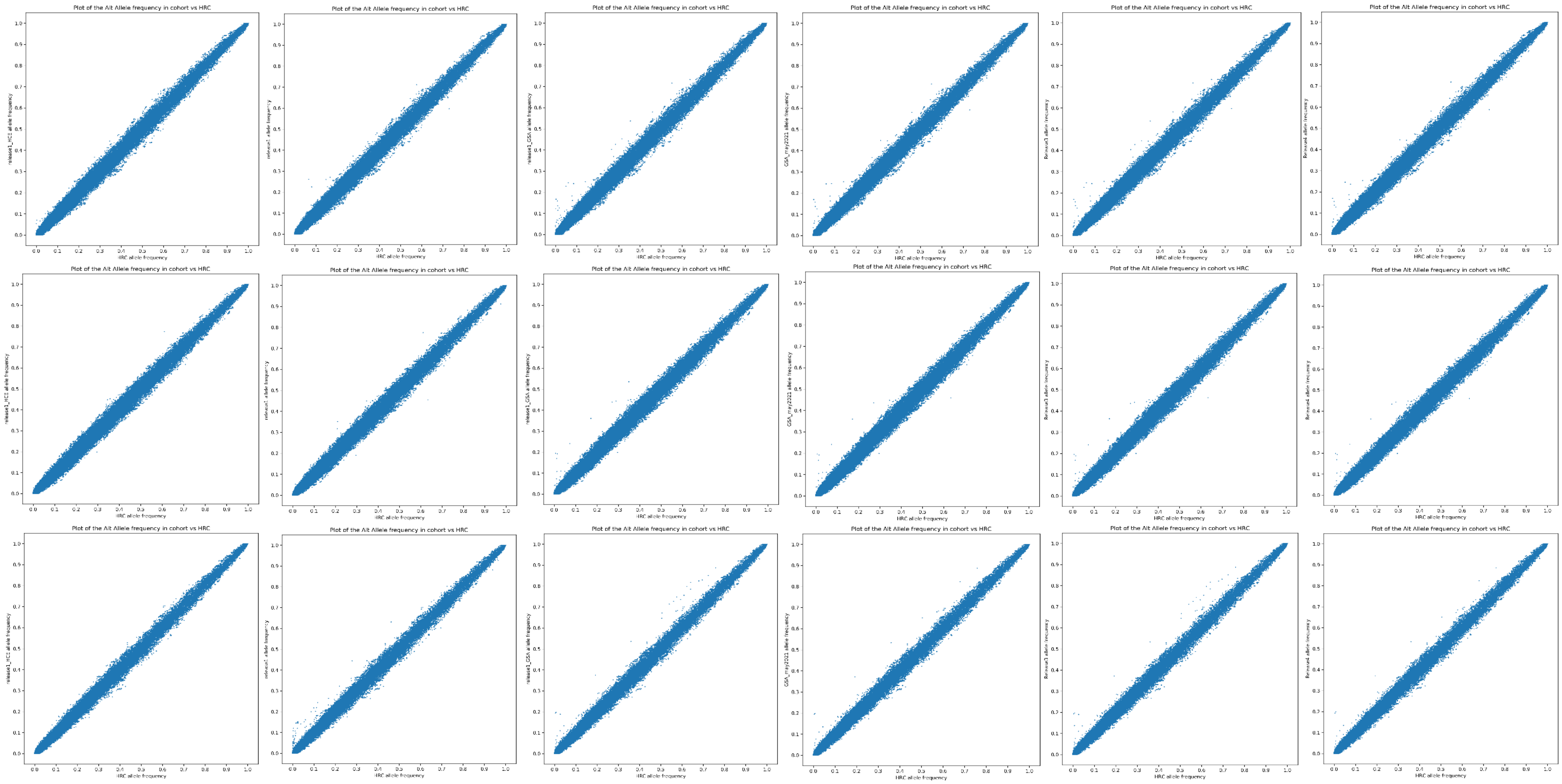

Plot of the Alt Allele frequency in cohort vs HRC
HRC allele frequency
release1 allele frequency
release1_GSA allele frequency
GSA_may2021 allele frequency
Release3 allele frequency
Release4 allele frequency

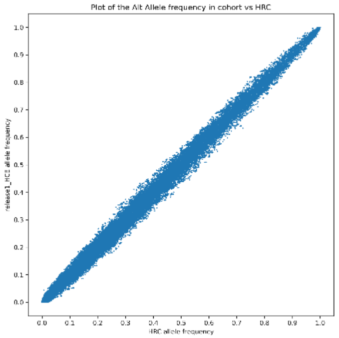
Plot of the Alt Allele frequency in cohort vs HRC
release1_HCE allele frequency
HRC allele frequency

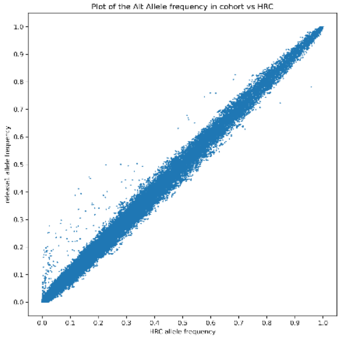
Plot of the Alt Allele frequency in cohort vs HRC
release1 allele frequency
HRC allele frequency

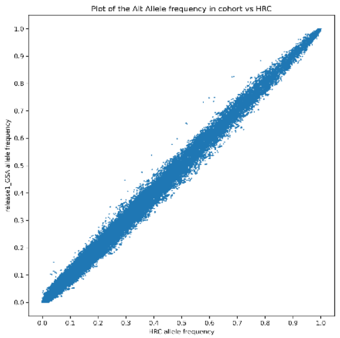
Plot of the Alt Allele frequency in cohort vs HRC
release1_GSA allele frequency
HRC allele frequency

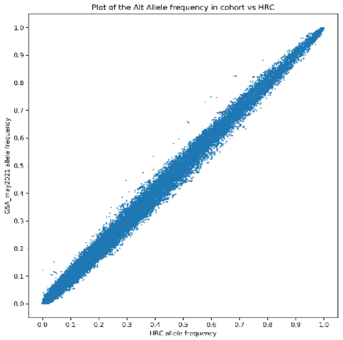
Plot of the Alt Allele frequency in cohort vs HRC
GSA_may2021 allele frequency
HRC allele frequency

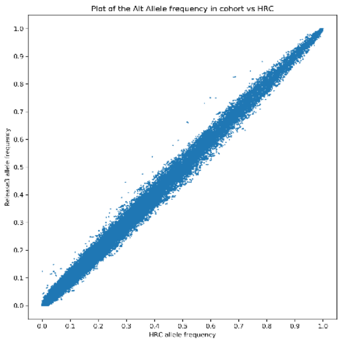
Plot of the Alt Allele frequency in cohort vs HRC
Release3 allele frequency
HRC allele frequency

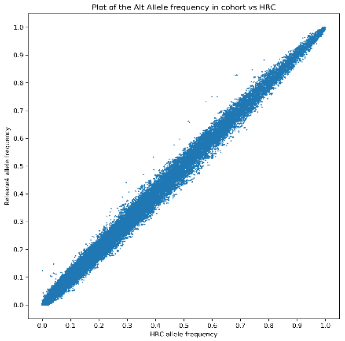
Plot of the Alt Allele frequency in cohort vs HRC
Release4 allele frequency
HRC allele frequency

Supplementary Figure 6. Imputation quality plots showing position vs imputation quality scores (INFO) for chromosome X and pseudoautosomal regions (PAR) and MoBa imputation batch. Plots from left to right are Imputation batch HCE, OMNI, GSA1, GSA2, GSA3, and GSA4. Plots from top to bottom are chromosome X, PAR1 and PAR2.

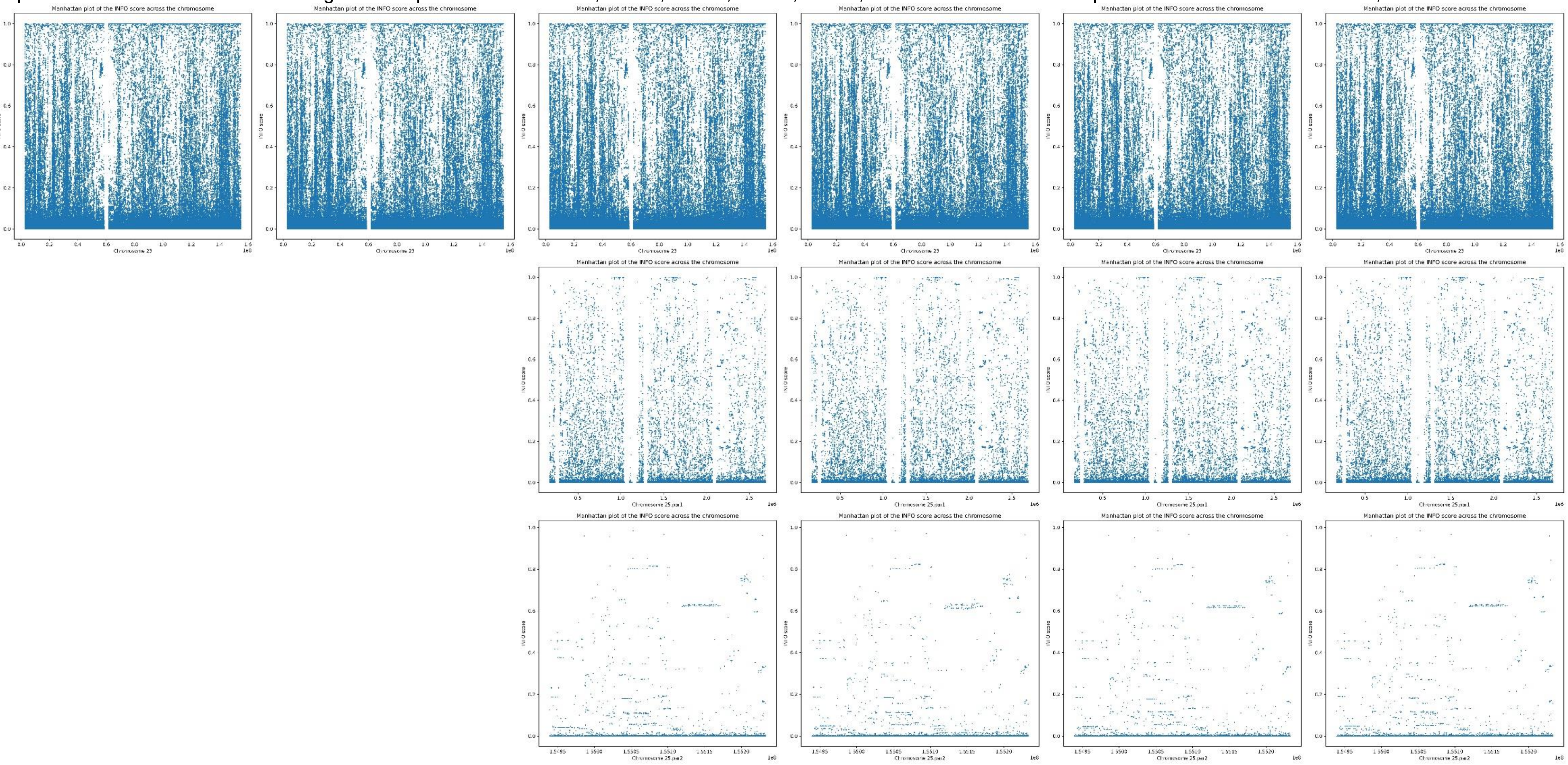

Supplementary Figure 7. Imputation quality plots showing minor allele frequency (MAF) in MoBa imputed data vs imputation quality scores (INFO) for chromosome X and pseudoautosomal regions (PAR) and imputation batch. Plots from left to right are Imputation batch HCE, OMNI, GSA1, GSA2, GSA3, and GSA4. Plots from top to bottom are chromosome X, PAR1 and PAR2.

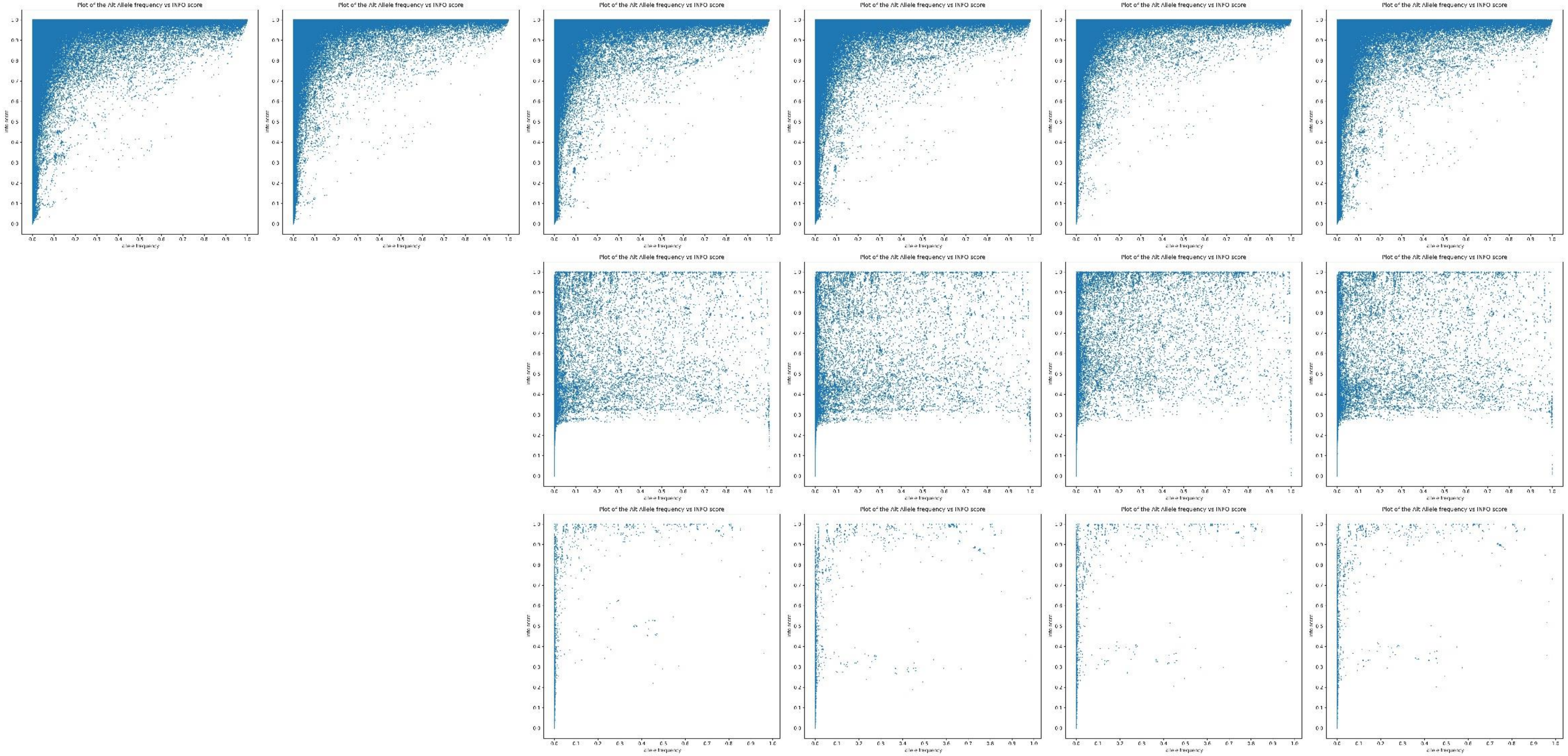

Supplementary Figure 8. Imputation quality plots showing minor allele frequency (MAF) in the Haplotype Reference Consortium vs MAF in MoBa imputed data for chromosome X and pseudoautosomal regions (PAR) and imputation batch. Plots from left to right are Imputation batch HCE, OMNI, GSA1, GSA2, GSA3, and GSA4. Plots from top to bottom are chromosome X, PAR1 and PAR2.

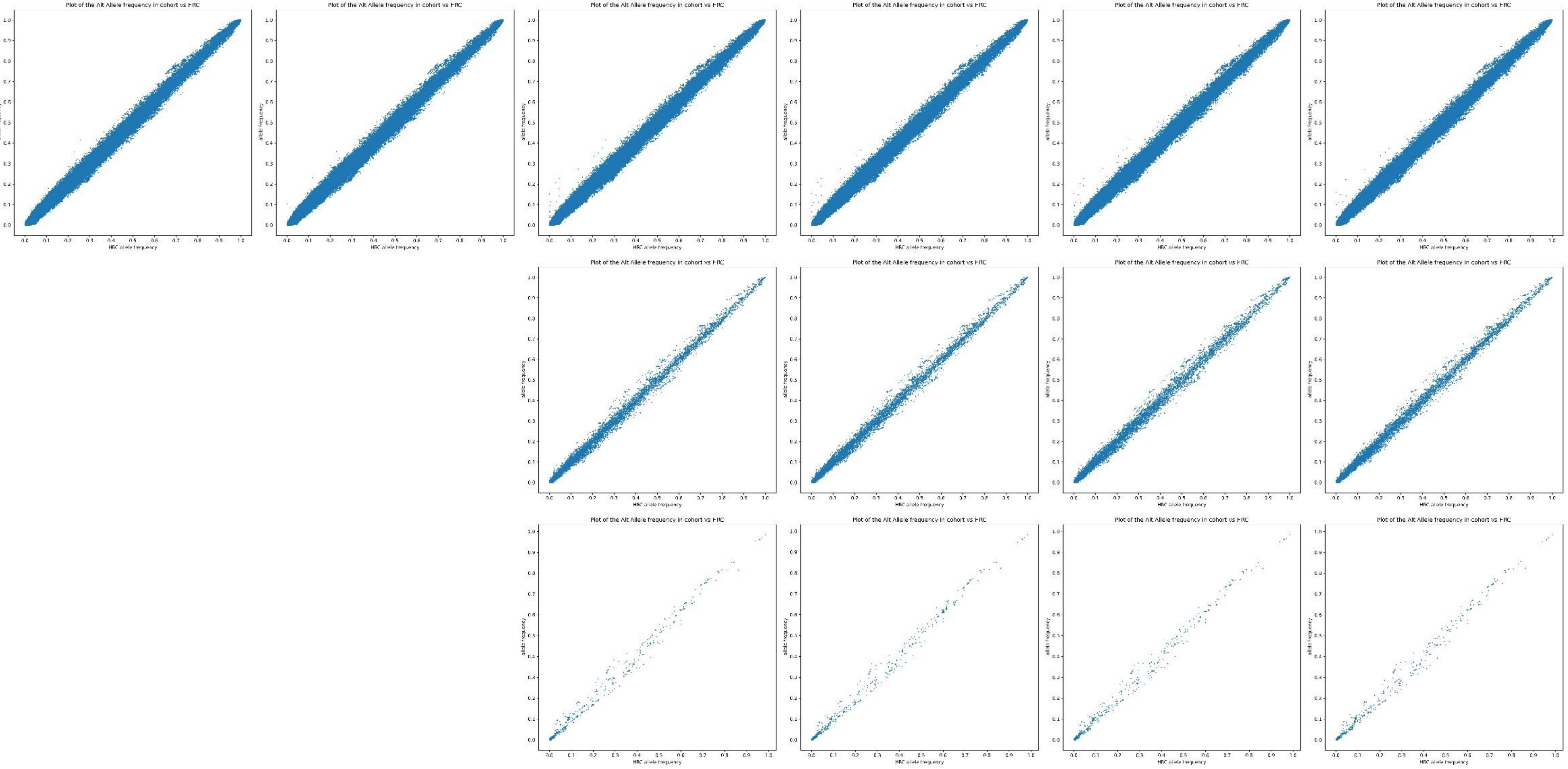
